# A novel biopolymer for mucosal adjuvant against respiratory pathogens

**DOI:** 10.1101/2022.09.07.506979

**Authors:** Ashley R. Hoover, Sunil More, Kaili Liu, Connor L. West, Trisha I. Valerio, Coline L. Furrer, Ningli Yu, Crystal Villalva, Amit Kumar, Lu Alleruzzo, Samuel S. K. Lam, Tomas Hode, Meng Zhao, James F. Papin, Wei R. Chen

## Abstract

Mucosal vaccinations for respiratory pathogens provide effective protection as they stimulate localized cellular and humoral immunities at the site of infection. Currently, the major limitation of intranasal vaccination is using effective adjuvants capable of withstanding the harsh environment imposed by the mucosa. Herein, we describe the efficacy of using a novel biopolymer, N-dihydrogalactochitosan (GC), as a nasal mucosal vaccine adjuvant against respiratory infections. Specifically, using COVID as an example, we mixed GC with recombinant SARS-CoV-2 trimeric spike (S) and nucleocapsid (NC) proteins to intranasally vaccinate K18-hACE2 transgenic mice, in comparison with Addavax (AV), an MF-59 equivalent. In contrast to AV, intranasal application of GC induces a robust, systemic antigen-specific antibody response and increases the number of T cells in the cervical lymph nodes. Moreover, GC+S+NC-vaccinated animals were largely resistant to the lethal SARS-CoV-2 challenge and experienced drastically reduced morbidity and mortality, with animal weights and behavior returning to normal 22 days post-infection. In contrast, animals intranasally vaccinated with AV+S+NC experienced severe weight loss, mortality, and respiratory distress, with none surviving beyond 6 days post-infection. Our findings demonstrate that GC can serve as a potent mucosal vaccine adjuvant against SARS-CoV-2 and potentially other respiratory viruses.

## Introduction

Worldwide, respiratory tract infections are the most common type of infection found in both adults and children^1^. They manifest a variety of symptoms generally limited to the upper respiratory tract, but occasionally they can become more sinister and result in pneumonia, significant lung damage, and/or even death, predominantly in young children or elderly adults^2^. Successful respiratory viral vaccination requires the development of a broad repertoire of antibodies (humoral immunity, IgA and IgG), T Helper 1 (Th1, interferon gamma: IFN-γ), and cytotoxic T cell responses (cellular immunity) systemically and specifically at the site of infection. Mucosal vaccination for respiratory viruses is an attractive option as it can trigger both humoral and cellular immunities locally and systemically^3–5^.

The current intranasal vaccines approved for human use, such as FluMist and NASOVAC, employ live attenuated virus (LAV)^6^. LAVs are attractive because they generate broader, more robust, and more durable immune responses against the targeted pathogen and possible variants^7^. Moreover, LAVs mimic natural infection and thus stimulate an effective immune response. Despite their effectiveness, there are drawbacks to LAVs. They are temperature sensitive and can regain virulence in some cases and are thus not recommended for individuals with severely compromised immune systems^8–10^. Additionally, during the attenuation process, the immunogenicity of the pathogen can become compromised, resulting in only moderate immune protection^6, 11^. Whole inactivated viruses and subunit vaccines do not have the issue of temperature sensitivity or the ability to regain virulence, but the immunogenicity of these vaccines can be compromised^10, 12, 13^. To increase the immunogenicity of inactivated viruses and subunit vaccines, adjuvants are needed to enhance the cellular and/or humoral immune responses to the targeted pathogens^14, 15^.

Development of adjuvants for mucosal vaccines faces significant challenges due to the nature of the mucosa, which contains several components that provide a protective barrier between the outside environment and our internal organ systems. It consists of a single epithelial layer that is covered with protective mucus, antimicrobial proteins, and symbiotic microbiota that maintain a low pH ^16, 17^. The epithelial layer is peppered with intraepithelial tissue-resident innate and adaptive immune cells, and specialized mucosal associated lymphoid structures capable of responding to pathogen invasion^16, 17^. Thus, an ideal mucosal adjuvant must be able to withstand the harsh environment of the mucosal surface, maintain the antigen at the site of inoculation for an extended period, activate/recruit immune cells to the site of administration, and stimulate an immune response like the naturally infecting pathogens. For respiratory viral pathogens, the ideal adjuvant needs to drive a type I interferon response, since these cytokines play a critical role in initiating antiviral immunity and generating tissue-resident immune cells in the respiratory epithelium^18, 19^.

Using adjuvants, such as cyclic dinucleotides, which elicit the production of type I IFN through STING signaling, leads to enhanced humoral and cellular immunities capable of preventing and eliminating intracellular pathogen infections following vaccination^20, 21^. However, cyclic dinucleotides must be administered repeatedly, in high quantities, and/or modified to prevent host cell inactivation^20–22^.

To overcome these and other limitations of existing mucosal adjuvants, we developed N-dihydrogalactochitosan (GC), a new molecule that is synthesized from galactose and chitosan^23^. Compared to the immune responses induced by its chitosan precursor^24, 25^, GC exhibits superior potency in inducing the type I IFN pathway activation^26^. In addition, GC has improved solubility over chitosan and retains many of its favorable properties, including its low toxicity, biodegradability, low-pH resistance, biocompatibility, and mucoadhesive nature.

Chitosan is currently used in a variety of chemical, agricultural, pharmacological, and medicinal applications^27, 28^. However, because of its poor solubility, high potential to be contaminated with endotoxins, and poor characterization of the polymeric mixture, chitosan has limited uses as a commercial medicinal product^29^. GC has been synthesized and purified under GMP conditions, with comprehensive structural characterization, extensive testing for metals, endotoxins, and other impurities, and with superior potency and improved solubility over its corresponding chitosan starting material, further addressing the major challenges in applications of chitosan. GC is currently developed as a medicinal product (referred to as IP-as a multi-modal immune stimulant to generate a systemic antitumor immune response when administered in conjunction with tumor ablation^30–32^.

In the present work, GC was used as a mucosal adjuvant for intranasal vaccines due to its functions as an immunostimulant/adjuvant. To mitigate the drawbacks of LAVs as mentioned, we used recombinant SARS-CoV-2 trimeric spike (S) and nucleocapsid (NC) proteins with GC for nasal vaccine formulation. K18-hACE2 mice were immunized intranasally with GC+S+NC and compared with Addavax (AV), an MF-

59 equivalent. We found that GC produces a robust, systemic antigen-specific antibody response and increases the number of T cells in the cervical lymph nodes following intranasal vaccination to a greater degree than AV. More importantly, animals vaccinated with GC+S+NC were strongly resistant to the lethal SARS-CoV-2 viral challenge and experienced drastically reduced weight loss and mortality, with complete recovery 22 days post-infection. In contrast, AV+S+NC-vaccinated animals all succumbed to infection within 6 days of the SARS-CoV-2 viral challenge, while experiencing severe weight loss, mortality, and respiratory distress. Our findings make a strong case for using GC as a mucosal adjuvant for intranasal vaccine. Not only does GC alone drive an antiviral response in antigen-presenting cells (APCs), but also simply combining it with recombinant proteins creates a highly effective mucosal vaccine against coronaviruses.

## Results

### GC retains antigen at the administration site and activates a variety of pattern recognition receptors involved in viral responses

Since one of the important functions of an effective vaccine adjuvant is to retain antigens at the administration site, we examined GC’s ability to retain ovalbumin labeled by Texas-Red (OVA-TR). We mixed GC of different concentrations (0.5, 0.75, 0.9, or 1%) with OVA-TR and injected the mixture subcutaneously in the flank of C57BL/6 mice, followed with whole-body fluorescent imaging over a period of seven days (Supplemental Figure 1A). We observed that GC at a higher concentration (0.9 and 1.0%) was able to retain OVA at the administration site significantly longer than PBS (Figure 1A, Supplemental Figure 1B). Specifically, ∼50% of the OVA was retained at the administration site by the 1% GC solution three days post-injection (Figure 1A).

**Figure 1.**
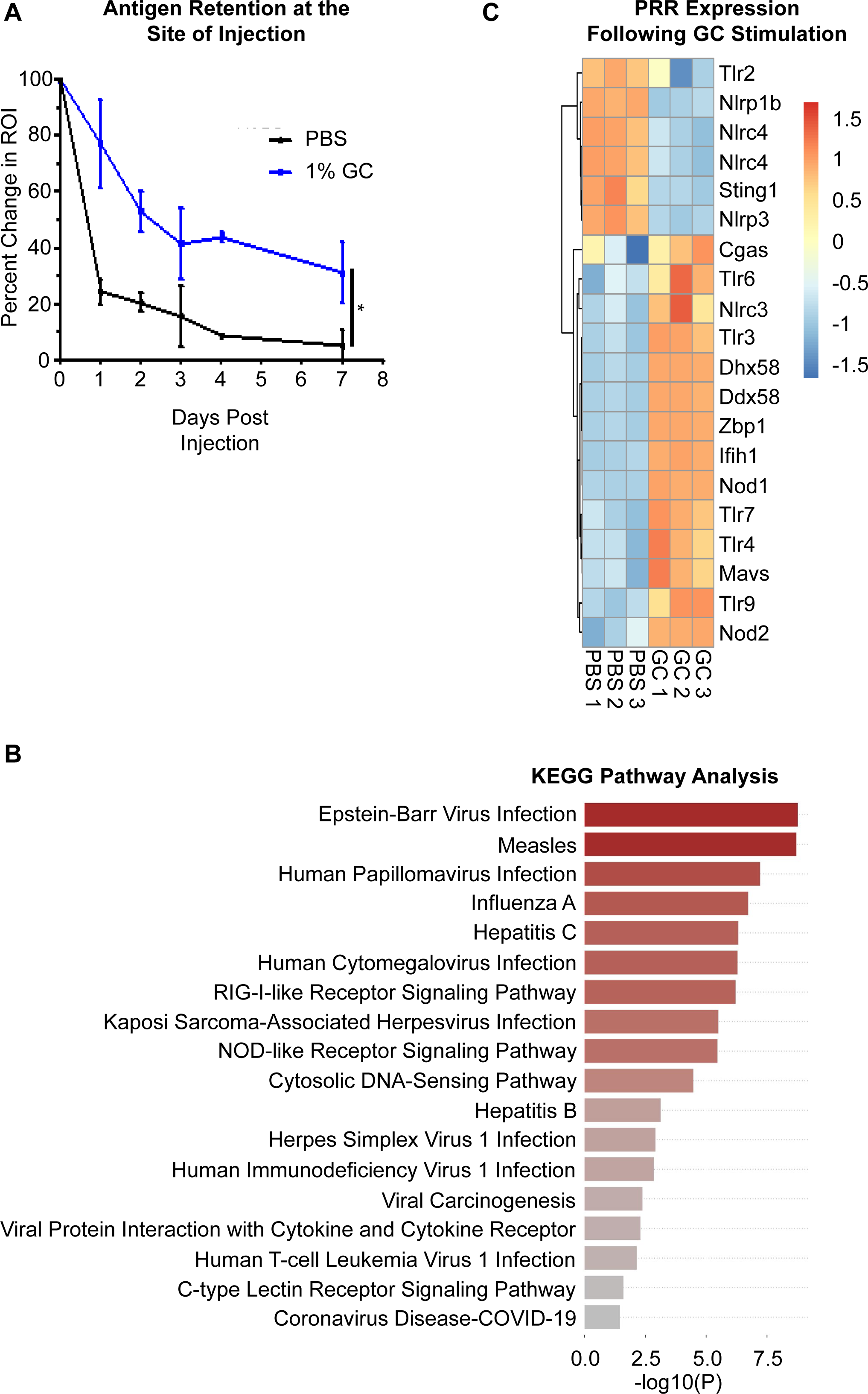
N-dihydrogalactochitosan retains antigen at the site of injection and stimulates a variety of antiviral pathways. A. Retention of antigen by GC. GC (1%) was mixed with ovalbumin labeled by Texas-Red (OVA-TR) and injected intradermally into the flank of C57BL/6 mice. Whole-body fluorescent imaging was performed over a period of several days. The percentage rate of change was calculated to quantify the amount of OVA localized at the injection site. Two-way ANOVA was used for statistical analysis. B. Stimulation of antiviral response pathways by GC. Wild-type BMDCs were stimulated +/- GC for 24 hours prior to harvesting for bulk RNA-sequencing analysis. KEGG Pathways Analysis revealed that GC stimulated a variety of antiviral response pathways. C. Heatmap showing the expression of various pattern recognition receptors (PRRs) in PBS- and GC-stimulated BMDCs (n=3).

Furthermore, when GC alone was delivered intranasally, at a concentration of 0.75%, the total CD45^+^ cellularity and B cell cellularity in the cervical lymph nodes (cLNs) were significantly increased 72 hours post-inoculation (Supplemental Figure 1C). In addition, we observed a trending increase in the number of migrating and resident dendritic cells (DCs), and CD4^+^ and CD8^+^ T cells in the cLNs (Supplemental Figure 1C). These data suggest that GC can induce a local response, leading to the migration and activation of lymphocytes in the nasal-associated lymphoid tissue (NALT).

Since we found that GC specifically activates innate immune cells, particularly DCs^26^, we decided to focus on the immune responses, particularly anti-viral responses, of bone-marrow-derived dendritic cells (BMDC) induced by GC. Specifically, to determine the changes in gene expression in GC-stimulated BMDCs, we performed bulk RNA-sequencing analysis. Our results revealed that a variety of gene expression pathways involved in anti-viral responses, including responses against Covid-19, were highly enriched in GC-stimulated BMDCs (Figure 1B). Several pathways involved in antigen processing and presentation, cytokine-cytokine receptor signaling, cell adhesion, cell-cell interaction, and antibody production networks, were also upregulated (Supplemental Figure 1D). Moreover, a variety of pattern recognition receptors (PRRs), most notably a diverse array of nucleic acid PRRs, including *Cgas, Zbp1, Tlr3, Tlr7, Tlr9, Mavs,* and *Ifih1,* were activated in BMDCs in response to GC (Figure 1C). Taken together, these results suggest that GC acts as a potent adjuvant for recombinant protein vaccination as it activates multiple PRRs, drives the expression of genes required for T cell activation and differentiation, and triggers a variety of viral response pathways required to initiate antiviral immunity.

### GC drives strong antigen-specific humoral immune responses following a two-dose vaccine regimen

To examine the adjuvant function of GC in inducing humoral immune responses, we prepared a recombinant viral protein (RVPs)-based vaccine, by mixing 1% GC with 5μg of either recombinant SARS-CoV-2 trimeric spike (S) protein or nucleocapsid (NC) protein. C57BL/6 mice (8 weeks old) were vaccinated intranasally (I.N.) once (1X) with GC, GC+S, and GC+NC. The serum, lungs, and cLNs were collected for analysis two weeks later (Supplemental Figure 2A). Total serum IgG and IgA were unchanged amongst all the vaccinated groups (Supplemental Figure 2B) and S-specific antibodies were not detected at this time point (data not shown). Lung cellularity and IgA levels were also unchanged in all the groups (data not shown). In the cLNs, total CD8^+^ T cells and naïve CD8^+^CD62L^+^ T cells were increased in the GC+NC 1X-vaccinated animals compared to PBS or GC groups (Supplemental Figure 2C). For CD4^+^ T cells, the memory CD44^+^CD62L^+^ CD4^+^ T cells were significantly increased after GC+NC vaccination compared to PBS controls (Supplemental Figure 2D), and B cell numbers were largely unchanged (Supplemental Figure 2E). While 1X vaccination was able to increase lymph node cellularity (Supplemental Figure 2C), it was unable to generate S-specific antibodies at a detectable concentration (data not shown). This suggests that one I.N. vaccination may not be sufficient to induce significant immune responses against viral infection.

We next vaccinated mice with S protein I.N. twice (2X), four weeks apart, and examined the humoral and cellular immune responses at different time points after vaccination (Figure 2A). We also used the MF-59 equivalent, Addavax (AV), as a comparing adjuvant, because MF-59 is currently used in human vaccines against viral infections (including influenza) and it has been demonstrated to enhance the mucosal immunogenicity of recombinant protein vaccination^14, 33^. PBS, 1% GC, or AV (15μl) was mixed with 2.5μg S protein and used for the two-dose vaccination scheme. Two weeks after the second vaccination, left lung lobe homogenates were acquired and total IgA and S-specific IgA were measured via antibody ELISA (Figure 2B). While not significant by One-Way ANOVA, total IgA in both GC+S- and AV+S-vaccinated animals was trending higher than in PBS controls. Only S-specific IgA was significantly higher in the GC+S-vaccinated animals compared to PBS controls (Figure 2B). Total serum IgG1 and IgG2c antibodies were measured one week after the first vaccination, one week after the second vaccination, and two weeks after the second vaccination (Figure 2C-D). One week after either the first vaccination or the second vaccination, there were no significant differences in total IgG1 among the four groups (Figure 2C). Two weeks after the second vaccination, total IgG1 in both GC+S and AV+S groups was trending up but not significantly by One-Way ANOVA (Figure 2C). Interestingly, S-specific antibodies were not detected in the PBS samples but were observed in both AV+S- and GC+S-vaccinated animals (Figure 2C). While IgG2c showed little change one week after the first vaccination, it is trending up in PBS+S-, GC+S-, and AV+S-vaccinated animals one week after the second vaccination (Figure 2D). Interestingly, two weeks after the second vaccination, IgG2c levels returned to the levels observed one week after the first vaccination (Figure 2D). Mirroring what was observed with S-specific IgG1 (Figure 2C), S-specific IgG2c was only represented in the AV+S- and GC+S-vaccinated groups, suggesting that GC drives a similar systemic antigen-specific antibody response to AV+S vaccination (Figure 2D). Taken together, these data indicate that GC induces a satisfactory antigen-specific humoral immune response that is comparable and possibly superior to a known vaccine adjuvant, AV, when intranasally delivered.

**Figure 2.**
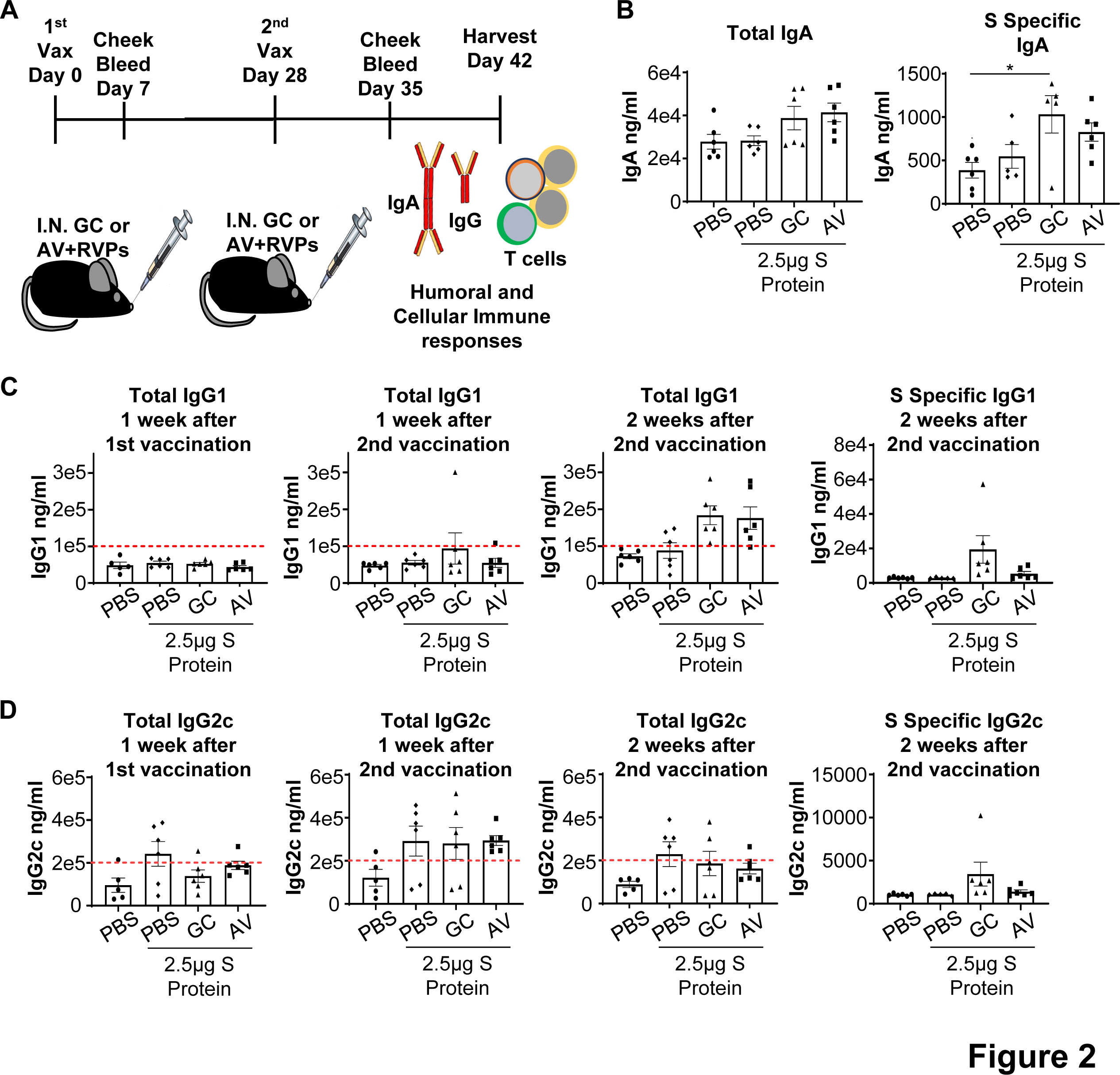
Multiple intranasal vaccinations using GC with recombinant SARS-CoV-2 spike protein drives localized, systemic antigen-specific humoral responses, in comparison with the adjuvant MF-59 equivalent, Addavax (AV) A. Schematic of the intranasal (I.N.) vaccination (2X) strategy and analysis using GC or AV as the adjuvant. B. Total IgA and S-specific IgA from left lung lobe homogenates harvested two weeks after the second vaccination. One-way ANOVA was used for statistical analysis. C. Total serum IgG1 and S-specific IgG1 levels one week after the first vaccination, one week after the second vaccination, and two weeks after the second vaccination. D. Total serum IgG2c and S-specific IgG2c levels one week after the first vaccination, one week after the second vaccination, and two weeks after the second vaccination.

### GC drives strong cellular responses following a two-dose vaccination regimen

In addition to the humoral immune responses, we examined the numbers of T and B cells residing within the cLNs and the lungs following the same vaccination schedule described in Figure 2A. While not significant by one-way ANOVA, there was a trending increase in the total CD8^+^ T cells and specific CD8^+^ T cell subsets in the cLNs of GC+S- or AV+S-vaccinated animals (Figure 3A). Interestingly, GC+S-vaccinated animals experienced a trending increase in CD8^+^CD44^+^CD62L^+^ and CD8^+^CD62L^+^ T cells while AV+S-vaccinated animals demonstrated a trending increase in CD8^+^CD44^+^ and CD8^+^CD62L^+^ T cells (Figure 3A). In contrast, CD4^+^CD44^+^ and CD4^+^CD44^+^CD62L^+^ T cells were both significantly increased in the cLNs of GC+S-vaccinated animals but not in the AV+S group (Figure 3B). B cells were not significantly affected in all experimental groups (Figure 3C).

**Figure 3.**
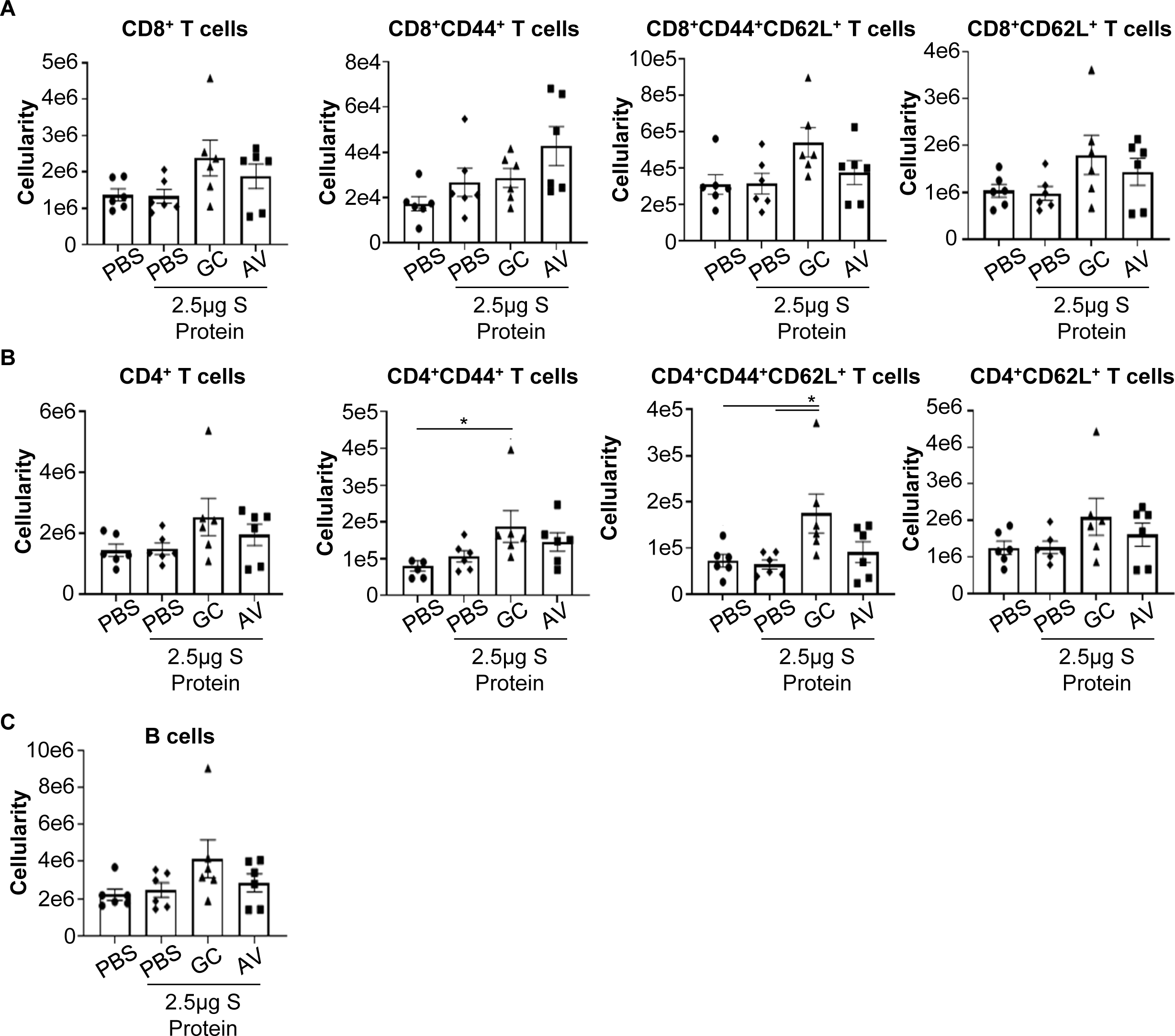
Intranasal administration of GC with recombinant SARS-CoV-2 spike protein increases cervical lymph node cellularity. A-C. Activation of leukocytes after two-dose vaccination. Cervical lymph nodes were isolated two weeks after the second intranasal vaccination and analyzed via flow cytometry. A. Activation of CD8^+^ T cells after two-dose vaccination. CD8^+^ T cells were gated on Live, CD45^+^CD3^+^CD8^+^ cells prior to analysis with CD44 and CD62L. Total cellularity of CD8 T cells, CD8^+^CD44^+^, CD8^+^CD44^+^CD62L^+^, and CD8^+^CD62L^+^ was calculated. One-way ANOVA was used for statistical analysis. B. Activation of CD4^+^ T cells after two-dose vaccination. CD4^+^ T cells were gated on Live, CD45^+^CD3^+^CD4^+^ cells prior to analysis with CD44 and CD62L. Total cellularity of CD4 T cells, CD4^+^CD44^+^, CD4^+^CD44^+^CD62L^+^, and CD4^+^CD62L^+^ was calculated. One-way ANOVA was used for statistical analysis. C. Activation of B cells after two-dose vaccination. B cells were gated on Live, CD45^+^CD19^+^B220^+^ cells and then calculated for total cellularity. One-way ANOVA was used for statistical analysis.

Next, we examined the lung cellularity following the two-dose vaccine regimen. GC+S-vaccinated animals did not experience a significant increase in CD8^+^ T cells, CD4^+^ T cells, or B cells (Supplemental Figure 3A-C). However, AV+S-vaccinated animals appeared to have a trending increase in CD8^+^ T cells and CD8^+^ T cell subsets (Supplemental Figure 3A). Total CD4^+^ T cells were trending up in AV+S-vaccinated animals, but this was not reflected in the three CD4^+^ T cell subsets examined (Supplemental Figure 3B). Furthermore, B cell numbers also had a trending increase in AV+S-vaccinated animals, which was not observed in the other three groups. Overall, the T cell cellularity within the lungs following a two-dose I.N. vaccination regimen resulted in minimal changes.

### Combining GC with the trimeric spike protein and the nucleocapsid protein increases the influx of T cells into the lung following intranasal vaccination

Next, to increase the cellular response in the lung following vaccination and to increase the coverage of antigens, we examined the combined effects of NC protein + S protein. The NC protein is one of the most abundant proteins in SARS-CoV-2, as it is required for genome replication and packaging, and it induces an immune response during the early stage of infection^34, 35^. To examine whether the addition of the NC protein could increase the robustness of recombinant protein vaccine responses, we vaccinated animals 2X, 4 weeks apart, with either GC+5μg S, GC+5μg NC, or GC+ 2.5μg S + 2.5μg NC. Two weeks after the second vaccination, the serum, lungs, and cLNs were examined for antibody and cellular responses. In the serum, we found that total IgG1 and IgG2c antibody levels were largely unchanged in the GC+S-, GC+NC-, or GC+S+NC-vaccinated mice compared to the PBS controls (Figure 4A-B). However, when examining S-specific IgG1 and IgG2c we found that GC+S+NC, with a reduced dose of S and NC proteins (2.5μg), induced equivalent S-specific IgG1 and IgG2c antibody levels compared to GC with a higher dose of S protein (5μg) (Figure 4A-B). A similar trend was observed in the lungs with total IgA and S-specific IgA (Figure 4C), with GC+S+NC producing significantly more S-specific IgA than PBS alone- or GC+NC-vaccinated animals.

**Figure 4.**
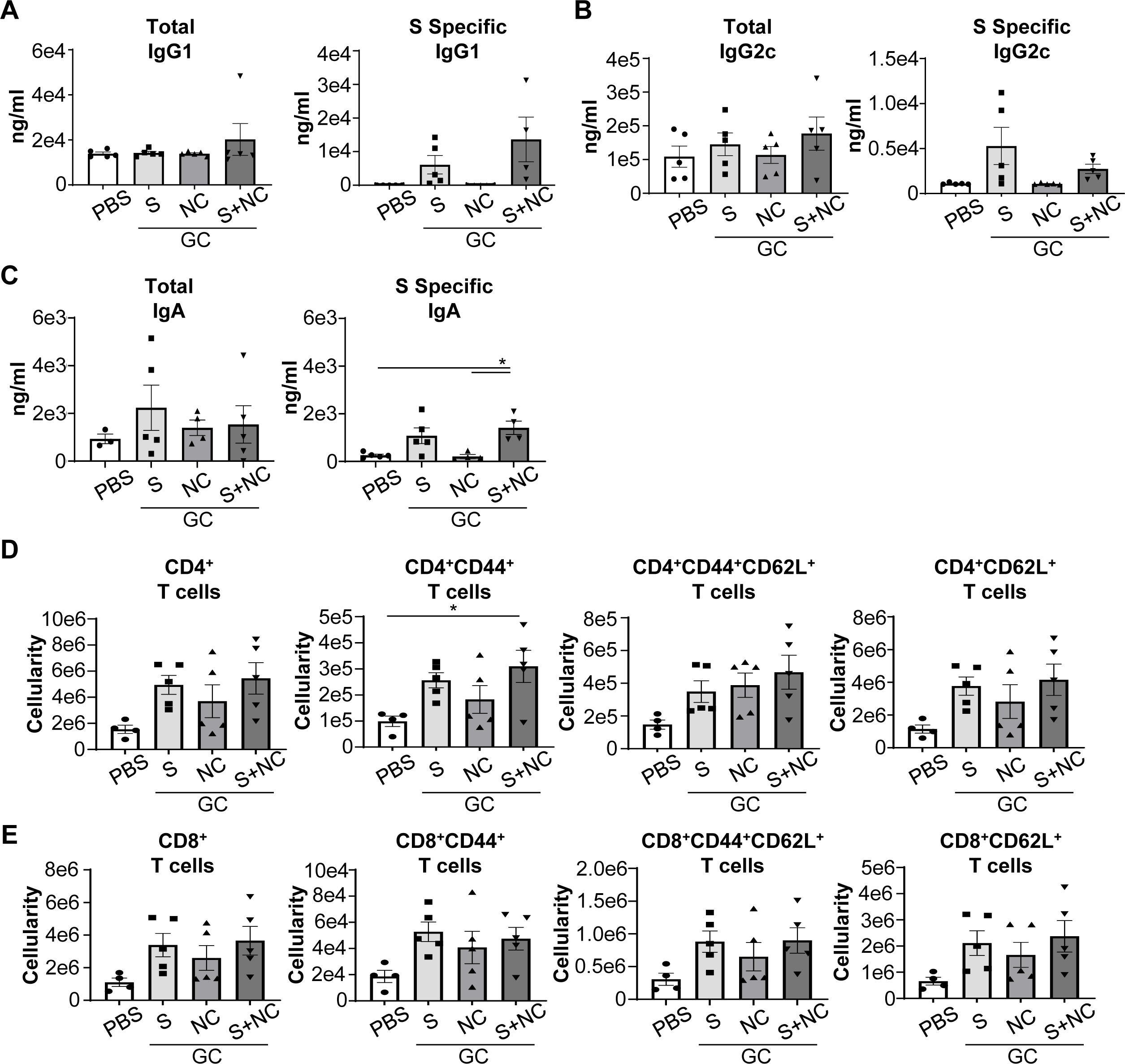
Combining recombinant SARS-CoV-2 spike and nucleocapsid proteins for intranasal vaccination enhances humoral and cellular responses. Antibody production and T cell activation were induced by intranasal vaccination via the combination of GC with different proteins: GC+NC (with 5 μg NC), GC+S (with 5 μg S), and GC+S+NC (with 2.5 μg S and 2.5 μg NC). A. Total Serum IgG1 and S-specific IgG1 levels two weeks after the second vaccination. One-way ANOVA was used for statistical analysis. B. Total Serum IgG2c and S-specific IgG2c levels two weeks after the second vaccination. One-way ANOVA was used for statistical analysis. C. Total IgA and S-specific IgA from left lung lobe homogenates harvested two weeks after second vaccination. One-way ANOVA was used for statistical analysis. D-E. T cell activation by GC with recombinant proteins. Cervical lymph nodes were isolated two weeks after the second intranasal vaccination and analyzed via flow cytometry. D. CD4^+^ T cells were gated on Live, CD45^+^CD3^+^CD4^+^ cells prior to analysis with CD44 and CD62L. Total cellularity of CD4 T cells, CD4^+^CD44^+^, CD4^+^CD44^+^CD62L^+^, and CD4^+^CD62L^+^ was calculated. One-way ANOVA was used for statistical analysis. E. CD8^+^ T cells were gated on Live, CD45^+^CD3^+^CD8^+^ cells prior to analysis with CD44 and CD62L. Total cellularity of CD8 T cells, CD8^+^CD44^+^, CD8^+^CD44^+^CD62L^+^, and CD8^+^CD62L^+^ was calculated. One-way ANOVA was used for statistical analysis.

In the cLNs of the GC+S-, GC+NC-, or GC+S+NC-vaccinated mice, trending increases in total and CD4^+^ and CD8^+^ T cell subsets (Figure 4D-E) were observed, with only the CD4^+^CD44^+^ T cells being significantly increased in GC+S+NC-vaccinated animals (Figure 4D). In the lungs, using either 5μg S protein or 5μg NC protein, instead of using 2.5μg S which was used for the experiments shown in Figures 2 and 3, resulted in a trending increase in total CD4^+^ and CD8^+^ T cells and their subsets and B cells (Supplemental Figure 4A-C). Combining 2.5μg S and 2.5μg NC proteins with GC resulted in an equivalent trending increase with 5μg S alone, suggesting that S and NC proteins are synergistic and can generate a more diverse immune response because it targets two separate SARS-CoV-2 proteins. For this reason, we chose to use the S+NC combination to vaccinate mice before viral challenge.

### Animals vaccinated by trimeric spike and nucleocapsid proteins + GC are resistant to lethal SARS-CoV-2 viral challenge

For viral challenge experiments, we vaccinated C57BL/6 transgenic mice expressing the human angiotensin 1-converting enzyme 2 (ACE2) receptor under control of the cytokeratin-18 (K18) gene promotor (K18-hACE2 mice). K18-hACE2 mice were vaccinated using the two-dose regimen described in Figure 2A. Six weeks after the second vaccination, animals were challenged with a lethal dose of SARS-CoV-2 and monitored for 21 days (Figure 5A). Efficacy following virus challenge was compared for six vaccine combinations: PBS, PBS+S+NC, GC, GC+S+NC, AV, or AV+S+NC. Within the first seven days following the lethal viral challenge, all mice in the PBS- and the GC-vaccinated groups either died or were humanely euthanized due to severe clinical signs or weight loss (Figure 5B-C). One of six animals in the AV group survived, while two of ten PBS+S+NC-vaccinated animals survived. Unexpectedly, all nine AV+S+NC-vaccinated animals either died or had to be humanely euthanized (Figure 5B-C). Excitingly, in the GC+S+NC-vaccinated group, 13 of the 15 vaccinated mice experienced minimal weight loss on the first and second days post-infection and survived without obvious complications (Figure 5B-C). Only two animals in the GC+S+NC group experienced weight loss and died within eight days of the viral challenge.

**Figure 5.**
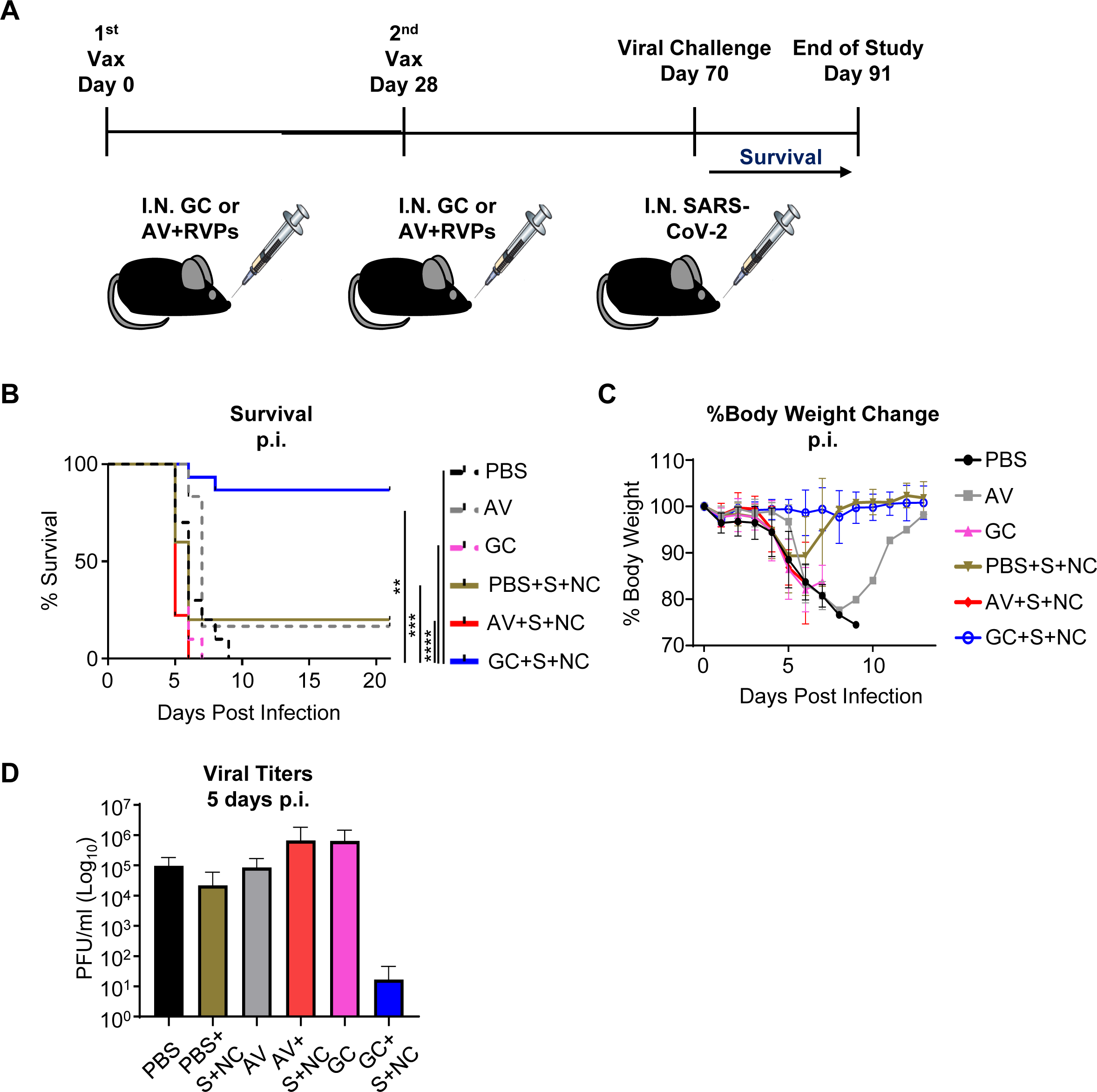
Intranasal vaccination with GC and recombinant SARS-CoV-2 spike and nucleocapsid proteins protects mice against lethal viral challenge. A. Schematic of the vaccination and viral challenge strategy for K18-hACE-2 animals. B. Survival rates of animals challenged by SARS-CoV-2 virus over a course of 21 days (n=6-15). C. Percent body weight change over a period of 21 days in virally infected animals. D. Plaque assays performed in lung homogenates on day 5 post viral infection (n=3 per group). Data represent mean ± SE of the respective groups. One-way ANOVA was used for statistical analysis.

As an alternative measure of vaccine efficacy to complement survival outcomes, we analyzed their viral loads in the lungs following infection. Five days post viral infection, three mice were randomly selected from each of the six groups and lung viral titers were assessed via plaque assays (Figure 5D). The mice in GC alone and AV+S+NC groups exhibited higher virus titers than that in the other groups, but the titers were not statistically significant with the number of animals used for analysis. Interestingly, only one mouse in the GC+S+NC group generated viral plaques, suggesting that GC+S+NC either mitigates virus replication in the lungs or reduces the amount of virus entering and infecting the lungs.

### GC + trimeric spike and nucleocapsid proteins protected animals from lung pathology following SARS-CoV-2 viral infection

The histopathology of the lungs from the six groups of intranasally vaccinated animals was analyzed at multiple time points post-infection. Prominent histopathology features were interstitial pneumonia, infiltration of inflammatory cells (lymphocytes, neutrophils, and macrophages), and bronchiectasia. The histology of the lungs and the pathology scores at day 0 are shown in Figures 6A and 6E. All the groups exhibited comparative pathology scores on day 3 (Figure 6B and 6F) when comparing a variety of lung pathology parameters. The most prominent features on day 3 were peribronchial and perivascular infiltration of lymphocytes (Figure 6B, empty circles). The interstitium was thickened and expanded with low numbers of lymphocytes, macrophages, and neutrophils (Figure 6B, arrow). On days 5 and 6 post-infection, the PBS, GC, and PBS+S+NC groups exhibited the most severe changes (Figure 6C and 6G-H). Bronchiectasia was evident on day 6 in the PBS, AV, and GC groups (Figure 6C, arrowhead). The bronchioles were infiltrated and expanded with moderate to large amounts of mucus admixed with inflammatory cells (bronchiectasia, Figure 6C, black arrows) in groups PBS+S+NC, AV+S+NC, and GC+S+NC. There were also moderate numbers of lymphocytes, neutrophils, and macrophages in the interstitium (Figure 6C, arrow) and extending into the alveoli (Figure 6C, empty arrowhead) in the three S+NC groups. In addition, the GC group exhibited moderate hemorrhages in the alveolar spaces (Fig. 6C; *). While the lungs of GC+S+NC-vaccinated mice exhibited peribronchiolar and perivascular inflammation six days after viral challenge, there was a complete absence of bronchiectasia (Figure 6C). While the lungs of GC+S+NC-vaccinated animals exhibited pathologic lesions, they were mild and were not to the level or severity experienced with the other groups. A mouse in the GC+S+NC-vaccinated group (Figure 6H) was sick and received high pathology scores but this was likely stochastic as survival rate in this group was significantly higher compared to the other groups. Lung pathology analysis 21 days post-infection revealed a significant reduction in the lung pathology in the GC+S+NC-vaccinated animals (Figures 6D and 6I). The GC+S+NC mice had characteristics of open alveolar spaces, and low amounts of perivascular and peribronchiolar cuffing as compared to AV and PBS+S+NC at day 21 (Figure 6D and 6I).

**Figure 6.**
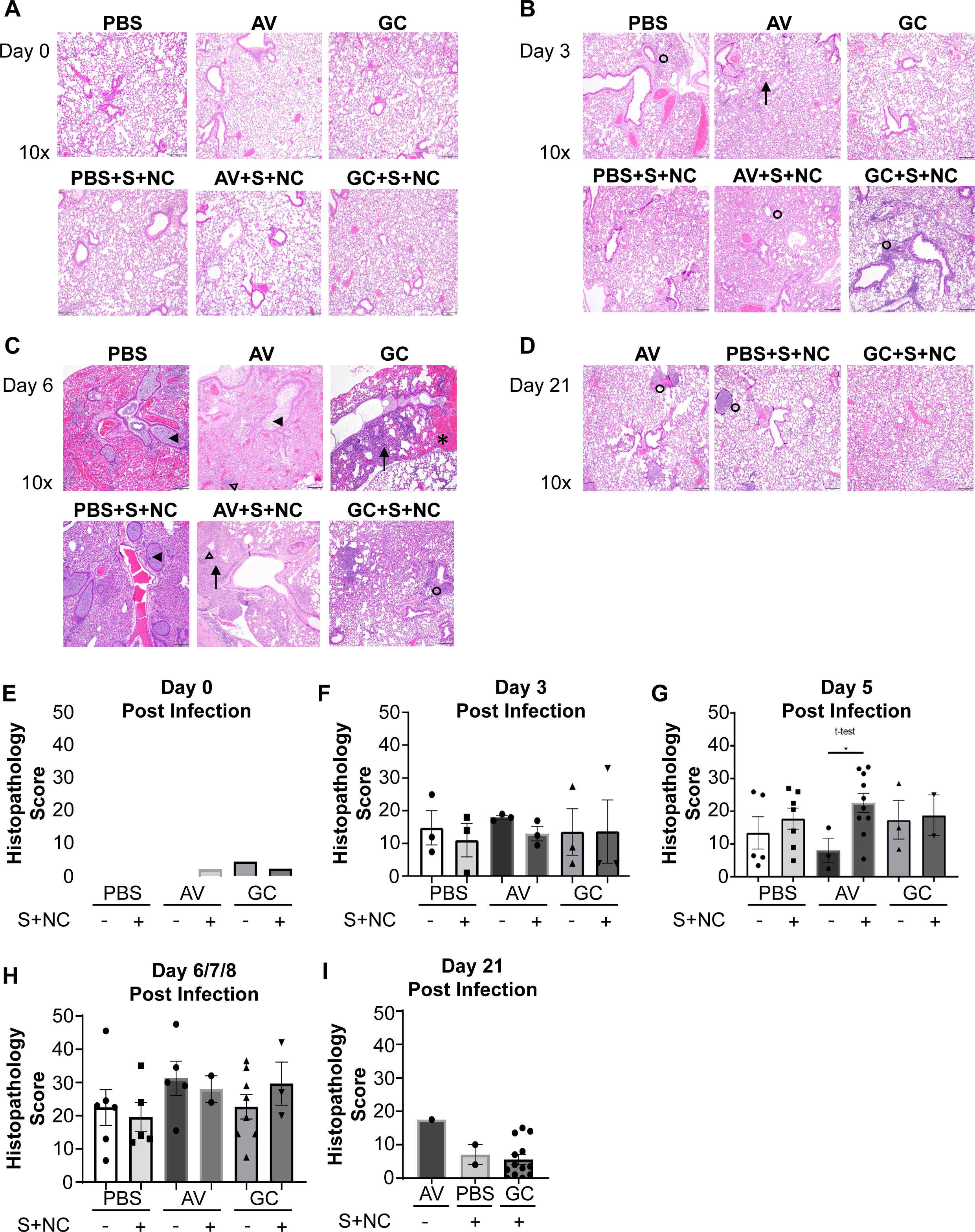
Intranasal vaccination with GC and recombinant SARS-CoV-2 spike and nucleocapsid proteins protects mice against severe lung pathology. A-D. 10x H&E sections of lungs at days 0, 3, 6, and 21 post SARS-CoV-2 challenge (n=6-15 per group). A. Day 0: H&E-stained lungs demonstrating normal lung architecture. B. Day 3: Interstitial (arrow) and peribronchial and perivascular immune infiltration (open circles) post-infection. C. Day 6: Bronchiectasia (arrowheads) and bronchioles infiltrated with mucus and immune cells (arrows) post-infection. Immune infiltration was also observed in the alveoli (empty arrowhead). Hemorrhages in the alveolar spaces (*) and peribronchial and perivascular immune infiltration (open circles) were observed. D. Day 21: Peribronchial and perivascular immune infiltration (open circles) post-infection. E-I. Pathological evaluation and scoring for the lung sections at days 0, 3, 5, 6/7/8, and 21 post-infection. The interstitium, alveoli, bronchioles, and vessels were evaluated for various parameters such as inflammation, edema, fibrin, necrosis, syncytial cells, hemorrhages, ectasia, vasculitis, and thrombi on a scale of 0-4: 0, no lesions; 1, minimal (1 to 10% of lobes affected); 2, mild (10 to 40%); 3, moderate (40-70%); and 4, severe (70 to 100%). The cumulative score was obtained from the 24 parameters examined. E. Cumulative histopathology scores for day 0. One-way ANOVA was used for statistical analysis. F. Cumulative histopathology scores for day 3. One-way ANOVA was used for statistical analysis. G. Cumulative histopathology scores for day 5. One-way ANOVA was used for statistical analysis. H. Cumulative histopathology scores for day 6/7/8 at the point of euthanasia. One-way ANOVA was used for statistical analysis. I. Cumulative histopathology scores for day 21. One-way ANOVA was used for statistical analysis.

### Effect of GC as an adjuvant in subcutaneous vaccination

To compare the effects of GC and AV as adjuvants for subcutaneous vaccination, GC+S+NC and AV+S+NC solutions were subcutaneously injected into the flank of the K18-hACE2 mice following the same two-dose regimen (Supplemental Figure 5A), with the same doses and same schedule, as for the intranasal vaccination. Six weeks after the second vaccination, animals were challenged with a lethal dose of SARS-CoV-2 and monitored for 21 days. As shown in Supplemental Figure 5B-C, only GC+S+NC and AV+S+NC protected the mice against the virus, with GC+S+NC resulting in a slightly higher survival rate than AV+S+NC.

## Discussion

Mucosal vaccination has the ability to induce local and systemic humoral and cellular immunities^3–5^, and mucosal adjuvants are critical to developing effective vaccines against respiratory viruses^36^. Key characteristics of successful vaccine adjuvants are their ability to enhance the immunogenicity of an antigen via retention and slow release, enhance the phagocytic capability of APCs, and reduce the amount of antigen required to stimulate protective immune responses^37^. This is reflected in their ability to boost humoral responses with antibody production and cellular responses through memory T cell formation. Moreover, vaccine adjuvants should drive an immune response similar to immune responses generated by the specific pathogen. Therefore, a type I IFN response would be ideal for respiratory viruses as these cytokines are required for antiviral immunity^38–40^. Thus far, most adjuvants currently used for human vaccines have been unable to activate a multitude of PRRs simultaneously; only one adjuvant, cytosine phosphoguanosine 1018 (CpG 1018), a TLR9 agonist, stimulates the production of type I IFN^14, 41, 42^. We developed N-dihydrogalactochitosan (GC), a novel biopolymer, as an effective mucosal vaccine adjuvant. In this study, we demonstrated GC’s adjuvant functions in a recombinant viral protein-based intranasal vaccine against respiratory pathogens, using SARS-CoV-2 as an example. Because of its physical and chemical properties, GC poses strong retention power in retaining antigens at the administration site at an appropriate concentration, as shown in Figure 1A and Supplemental Figure 1B. Furthermore, GC activated several different nucleic acid PRRs (Figure 1C), all of which were downstream of STING signaling and led to type I IFN and IL-1β production^26^. GC also activated immune pathways involved in response to Covid-19, as shown in Figure 1B.

In our previous work, we discovered that GC was able to produce a strong type I IFN response in tumor-infiltrating leukocytes (TILs) following its direct injection into the tumor^31, 43^. However, we also found that GC did not cause direct, non-specific activation of T cells, but instead works through the activation of APCs (data not shown). To determine the critical roles of DCs in antigen presentation, T cell activation, and differentiation, and effector and memory T cell formation, we discovered the pathways of T cell activation through DC activation, consistent with previous published results^44, 45^. We have previously shown that GC directly induces the maturation of DCs via increased co-stimulatory molecules CD80, CD40, and MHC-II^46^. Similarly, activation of DCs by GC was also observed using mouse DC cell line DC2.4^47^ or DCs isolated from a GC-injected tumor and tumor-draining lymph node^46^. In a follow-up study using murine BMDCs, it was found that GC activates DCs in part via STING signaling leading to IFNβ/α (type I IFNs) and IL-1β production^26, 44^.

We demonstrated that GC was also capable of activating multiple PRRs to induce the production of type I IFN and IL-1β, including nucleic acid sensing pathways that play a significant role in antiviral immunity (Figure 1C). Type I IFNs have been shown to provide a “third signal” to T cells responding to a presented antigen, enhancing their effector function and clonal expansion^48^. IL-1β enhances the expansion, differentiation, memory formation, and migration of antigen-primed CD4^+^ and CD8^+^ T cells ^49^. Moreover, the addition of IL-1β enhances the efficacy of weakly immunogenic vaccines^49^. We found that GC+S vaccination could recruit T cells into the axillary lymph nodes to a greater extent than AV+S (Figure 3). Furthermore, GC vaccination was able to significantly increase the cellularity of CD4^+^ T cells in the cervical lymph nodes (Figure 3B), but with the S protein alone, no changes were observed in the lung (Supplemental Figure 3A-B).

NC protein is highly immunogenic, highly conserved with a much slower mutation rate than the S protein, and drives robust T and B cell responses^50–53^. Therefore, to further enhance the influx of immune cells into the lung following intranasal vaccination, we combined the S protein with the NC protein. GC+S+NC generated S-specific antibody responses in the serum and lungs after two vaccinations (Figure 4A-C). More importantly, GC+S+NC increased the cellularity of T cells in the cervical lymph nodes and resulted in a trending increase of T and B cells within the lungs of the twice-vaccinated animals (Figure 4D-E, Supplemental Figure 4A-B). These results prompted us to vaccinate K18-hACE2 transgenic mice using S+NC mixed with either PBS, AV, or GC, followed by SARS-CoV-2 viral challenge with a lethal viral dose.

Six weeks after the second vaccination, K18-hACE2 transgenic mice were intranasally challenged with the SARS-CoV-2 virus. GC+S+NC vaccination potently prevented robust infection, morbidity (clinical signs and weight loss), and mortality, and exhibited significantly reduced viral load, and severe lung pathology, in response to the SARS-CoV-2 viral challenge (Figures 5 and 6). Surprisingly, AV+S+NC vaccination neither prevented mortality, nor reduced viral load, lung pathology, or weight loss. In fact, this group of animals performed significantly worse than all four control groups (PBS, GC, AV, PBS+S+NC) as shown in Figure 5. Lung pathology revealed that the AV+S+NC-vaccinated animals experienced severe bronchiectasia and interstitial inflammation 3 and 6 days post-infection (Figure 6B-C). This result was unexpected as the AV+S-vaccinated animals generated comparable humoral and cellular responses to GC+S in the lungs and cervical lymph nodes (Figures 2 and 3). These results suggest that only appropriate adjuvants should be used for lower respiratory tract infections and that the wrong type of adjuvants can make the lung pathology significantly worse.

Bronchial changes such as bronchiectasia are detrimental to COVID-19 patients^54^ as well as in a variety of other chronic lung diseases such as COPD^55^ and can also be seen in patients in the acute phase of SARS-CoV-2 infection^56^. We observed a tremendous increase in intraluminal bronchiolar mucus with inflammatory cells leading to the expansion of the bronchioles in all the control groups (Figure 6B-D). Interestingly, GC+S+NC-vaccinated mice did not exhibit this phenomenon, indicative of the lung protection by GC+S+NC against bronchiolar structural damage caused by the viral challenge (Figure 6B-D). Furthermore, the absence of bronchiectasia correlates well with the observation of the GC+S+NC-vaccinated mice experiencing little to no respiratory distress. These animals were likely well-ventilated, resulting in significantly reduced morbidity and mortality even though their cumulative histopathology scores were not significantly lower.

As mucosal adjuvants, GC and AV induced the same levels of humoral immune responses, as shown in Figure 2. The results of our protein-based vaccines using mucosal adjuvants GC and AV are not directly related to antibody production. Furthermore, neutralizing antibody titer results (data not shown) did not show significant differences between serum samples from different vaccinated groups. However, GC is proven to be more effective in protecting intranasally vaccinated mice than AV, as shown in Figure 5. These results indeed confirmed our hypothesis that GC+S+NC stimulates a strong cellular response which is not observed in the other groups. This could also help explain why the animals have lung pathology, indicating that the mice are becoming infected, but are able to clear the infection because of their adaptive T cell responses (Figure 6). In future studies, we will fully characterize the T cell responses responsible for keeping animals alive after lethal viral infections.

When GC and AV were used in sub-Q vaccination in combination with S and NC proteins, both adjuvants showed similar protecting abilities against the challenges with a lethal dose of SARS-CoV-2 virus, as shown by Supplemental Figure 5. GC is an effective mucosal adjuvant for intranasal vaccination because it induces the production of both type I IFN and IL-1β^26^. The in-depth mechanism of GC’s functions as a mucosal adjuvant for intranasal vaccination against respiratory pathogens will be investigated in future studies.

In addition, various adjuvants have been used, such as AS03 and AS04, for vaccines against respiratory pathogens ^57^. While these adjuvants do not induce immune responses in the same way GC does, future studies are warranted to compare the effects of AS03, AS04, and other adjuvants, which have been used in humans, with GC, particularly as mucosal adjuvants for intranasal vaccination.

Many IN vaccines that worked in mice failed in large animals, leading to limited choices of IN vaccines for humans. We are cautiously optimistic about GC, as there is no other mucosal adjuvant like it on the market for IN vaccination. In addition, GC has been used in humans for cancer treatment^27, 28, 30^. However, future pre-clinical studies and clinical trials are needed to determine the efficacy of GC + recombinant protein-based IN vaccines against COVID and other respiratory viruses.

In summary, the results presented herein reveal that GC acts as a potent mucosal vaccine adjuvant when administered in combination with recombinant viral proteins to prevent severe lung pathology and mortality in mice after SARS-CoV-2 infection. GC is effective as a viral vaccine adjuvant because it can retain the antigens in the mucosa for an extended period and it activates the same pathways involved in antiviral responses, thus driving potent antiviral immunity. Future work will focus on the understanding of GC’s localized immune effects on the nasal mucosa, longevity of the immune response, and altering parameters of the vaccination regimen to create the most effective intranasal immune response.

## Materials and Methods

### Animals

All animal studies were approved by the IACUC in Oklahoma Medical Research Foundation, the University of Oklahoma, and Oklahoma State University. C57BL/6 were purchased from Jackson Laboratories (Stock number: 000664). Wild-type C57BL/6 mice were used in this study to analyze the cellular and humoral responses to vaccination using either GC (Immunophotonics, Inc., St. Louis, MO) and/or Addavax (InvivoGen, cat# vac-adx-10). B6.Cg-Tg(K18-ACE2)2Prlmn/J were purchased from Jackson Laboratories (Stock number: 034860)^58^ and were used for vaccination and viral challenge with SARS-CoV-2 to test the efficacy of recombinant protein/adjuvant intranasal vaccination.

### Antigen Retention by GC

Soluble OVA labeled by Texas-Red (OVA-TR) was purchased from Life Technologies/Invitrogen (5mg in lyophilized powder) and reconstituted with 1.25ml PBS to achieve a concentration of 4mg/ml. OVA-TR was further diluted with PBS to a concentration of 0.8mg/ml and then mixed 1:1 with 2% IP-001, or PBS as control, to achieve a final solution with 0.4mg/ml OVA-TR in 1% GC. GC+OVA-TR solutions were also prepared with different GC concentrations (0.9, 0.75, and 0.5%). Then GC+OVA-TR of 100 μl volume was injected subcutaneously into the flank/back of shaved C57BL/6 mice. Whole-body fluorescent imaging (IVIS 100) was performed over a period of several days. A region of interest (ROI) was drawn at the injection site for quantification of fluorescence from Texas red, which serves as the surrogate for the amount of OVA. The percentage change in fluorescence over time was calculated by comparison to the starting value taken shortly after injection. Two-way ANOVA was used for statistical analysis.

### ELISA

Total IgA (Cat# 88-50450-88), IgG1 (Cat# 88-50410-86), and IgG2c (Cat# 88-50670-22) ELISA kits were purchased from ThermoFisher Scientific and were used according to manufacturer’s instructions for the serum samples. Lung samples were diluted 1/1000 for analysis. For analysis of Trimeric Spike specific antibodies, Nunc MaxiSorp ELISA plates (Biolegend, Cat# 423501) were coated with 10μg/ml trimeric spike protein overnight at 4°C. Serum samples were diluted 1/1000 while lung samples were diluted 1/100. After incubation, the plates were developed using the reagents provided in the Total Ig ELISA kits. Standards for total IgA, IgG1, and IgG2c were used to generate standard curves and calculate the concentration of spike-specific antibodies. Plates were read using the Bioteck Synergy H1 hybrid plate reader. Samples were analyzed and graphed using the GraphPad Prism Software. One-way AVOVA was used to determine the statistical significance between the groups being compared.

### Intranasal Vaccination and Immune Cell Isolation

Recombinant SARS-CoV-2 proteins were purchased from R&D systems. For vaccination, either 5μg or 2.5μg of the S protein or NC protein (R&D Systems, cat# 10474-CV-050 and cat# 10549-CV-100), both resuspended in PBS, were mixed with 15μl of 1% GC or 15μl of Addavax. The final volume delivered intranasally was 20μl, 10μl per nostril, delivered to male and female animals in equal numbers. Animals were vaccinated once or twice (four weeks apart). Two weeks after the first or second vaccination, the cervical lymph nodes, lungs, serum, and spleen were isolated. The serum was isolated via cardiac puncture and acid-citrate-dextrose (ACD) solution was added to prevent coagulation. The left lobe was used for antibody isolation, while the remaining lobes were used for flow cytometry. Lungs were digested using HEPES buffer containing LiberaseTM 20μg/ml (Roche, cat# 5401127001) and DNase I (100μg/ml) (Roche, cat # 4536282001) for 25 minutes at 37°C to isolate total leukocytes. Red blood cells were lysed using ACK lysis buffer and then resuspended in RPMI containing 10% FCS and Penicillin/Streptomycin until used for flow cytometry. Spleens and cervical lymph nodes were isolated and enzymatically digested in serum-free RPMI containing 100μg/ml Collagenase IV (Gibco, cat# 17104019) and 20μg/ml DNase I (Roche, cat# 4536282001) for 20 minutes at 37°C. After digestion, cells were washed in 1x HBSS, 5% fetal calf serum (FCS), and 5mM EDTA, and then for the spleens, the red blood cells were lysed using ACK lysis buffer. The cells were then resuspended in RPMI containing 10% FCS and Penicillin/Streptomycin until used for flow cytometry.

### Flow Cytometry

The isolated leukocytes from the lungs, cervical lymph nodes, and spleens were stained with the following antibodies: CD45-BV421 (Biolegend, Cat# 10314), CD40-Pacific Blue (Biolegend, Cat# 124626), Ghost Dye-Violet-510 (Tonbo Biosciences, Cat# 13-0870), CD11b-BV570 (Biolegend, Cat# 101233), CD103-APC (Biolegend, Cat# 121414), CD11c-AF647 (Biolegend, Cat# 117312), MHCII-RedFluor710 (Tonbo Biosciences, Cat# 80-5321), CD86-BV605 (Biolegend, Cat# 105037), Ly6G-SB702 (Invitrogen, Cat# 67-9668-82), Ly6C-BV785 (Biolegend, Cat# 128041), CD64-PE (Biolegend, Cat# 139304), F4/80 PE/Dazzle (Biolegend, Cat# 123146), B220-PE-Cy5 (Biolegend, Cat# 103210), CD19-PE-Cy7 (Invitrogen, Cat# 25-0193-82), CD24-FITC (Biolegend, Cat# 101806), CD44-PerCP-Cy5.5 (Tonbo Biosciences, Cat# 65-0441), CD8-BUV395 (BD Horizon, Cat# 565968), CD3-BUV496 (BD Optibuild, Cat# 741117), CD4-BUV563 (BD Horizon, Cat# 612923), CD62L-BUV737 (BD Horizon, Cat# 612833). Briefly, 5×10^6^ cells were stained with antibodies and placed on ice for 20 minutes in FACs buffer. Cells were washed and then fixed with Fixation Buffer (BD, Cat# 554655) according to the manufacturer’s instructions. The cells were then washed and resuspended in 1x PBS and then analyzed on the Cytek Aurora spectral flow cytometer. Flow cytometry data was then analyzed using FlowJo software version 10.7 and then graphed using GraphPad Prism. One-way AVOVA was used to determine the statistical significance between the groups being compared.

### Lung homogenate

At the time of isolation, the left lung lobe was isolated and snap-frozen in liquid nitrogen and stored at -80°C until use. To generate lung homogenate for ELISA, lung tissues were thawed on ice and then resuspended in 0.5ml of Tissue Protein Extraction Reagent (T-PER) (ThermoScientific, cat# 78510) containing Complete Mini Protease Inhibitor Cocktail (Roche, cat# 11836170001). Tissues were homogenized using an electric handheld tissue homogenizer and were stored at -80°C until used for ELISA.

### Viral Challenge

B6.Cg-Tg(K18-ACE2)2Prlmn/J were used for viral challenge experiments. The animals were bred in-house according to IACUC-approved protocols. Male and female mice heterozygous for the K18-hACE2 transgene were used for vaccination and viral challenge experiments. Animals were vaccinated as described above. Briefly, mice were vaccinated twice, four weeks apart. Six weeks after the second vaccination, the animals were challenged with a lethal dose of SARS-CoV-2 (isolate USA-WA1/2020, cat #52281) which was obtained from BEI Resources. The mice were anesthetized with Xylazine/ketamine anesthesia and inoculated intranasally with 2 × 10^4^ pfu/mice in 50 μl PBS (25 μl per nostril). Mice were monitored for clinical signs and weight changes every day. Representative mice were sacrificed at day 0 as controls and at days 3 and 5-7 post-infection. The mice that were moribund and or lost >25% body weight were humanely euthanized. The mice that survived the virus challenge were sacrificed on day 22 post-infection. Necropsy on each mouse was performed and tissues were harvested. The left lung was used for virus titer and the right lungs were inflated with 200 μl of 10% formalin for histopathology.

### Histopathology Scoring

Lungs from the infected and uninfected mice were infused with 200µl of 10% formalin and placed in a 10% formalin container. Lungs were trimmed, processed, and embedded in paraffin and 5μm sections were cut and stained with hematoxylin and eosin (H&E). The lung sections were scored blindly by two board-certified veterinary pathologists and averages were obtained for each mouse lung. The interstitium, alveoli, bronchioles, and vessels were evaluated for various parameters such as inflammation, edema, fibrin, necrosis, syncytial cells, hemorrhages, ectasia, vasculitis, and thrombi on a scale of 0-4: 0, no lesions; 1, minimal (1 to 10% of lobes affected); 2, mild (10 to 40%); 3, moderate (40-70%); and 4, severe (70 to 100%). The cumulative score from the 24 parameters examined was used for comparison using GraphPad Prism.

### Virus Titers

Left lung lobes were homogenized in 100 μl serum-free Opti-MEM (Gibco cat# 31985070) using a bead mix (Fisherbrand cat#15340153) in a tissue homogenizer (Fisherbrand cat#15340164). The homogenate was clarified with centrifugation at 2000 rpm x 5 mins. Fifty microliters of supernatant were serially diluted and 100 μl were plated on Vero cells (ATCC cat# CRL1586) monolayer in 12 well plates. The plates were incubated at 37°C for 1 hour with intermittent gentle shaking every 10 mins. After 1 hour post-infection, the media was removed and replaced with post-inoculation media containing overlayed with 0.6% Avicel (Dupont cat# CL-611NF) solution supplemented with 3% FBS (Gibco cat# 10437-28) and 2X MEM (Gibco cat# 11095-080). Three days post-infection overlay was removed, and cells were fixed with 10% neutral buffered formalin for 10-15 mins. The cells were then stained with 1% crystal violet solution (Sigma cat# HT90132) and plaques were counted for each dilution. The virus titers were determined by multiplying the number of plaques with the dilution factor.

### Subcutaneous vaccination and viral challenge

Either 5μg or 2.5μg of the S protein or NC protein, both resuspended in PBS, were mixed with 15μl of 1% GC or 15μl of Addavax. The final volume delivered subcutaneously was 20μl, for male and female animals in equal numbers. Animals were vaccinated twice (four weeks apart). Six weeks after the second vaccination, the animals were challenged with a lethal dose of SARS-CoV-2. The mice were anesthetized with Xylazine/ketamine anesthesia and inoculated intranasally with 2 x 10^4^ pfu/mice in 50 μl PBS (25 μl per nostril). Mice were monitored for survival and weight changes every day for 21 days. The vaccination regimen is shown in Supplemental Figure 5A.

## Acknowledgements

This work was supported in part by Immunophotonics, Inc. The following reagent was deposited by the Centers for Disease Control and Prevention and obtained through BEI Resources, NIAID, NIH: SARS-Related Coronavirus 2, Isolate USA-WA1/2020, NR-52281.

## Literature Cited

1. Gaunt, E.R., Harvala, H., McIntyre, C., Templeton, K.E. & Simmonds, P. Disease burden of the most commonly detected respiratory viruses in hospitalized patients calculated using the disability adjusted life year (DALY) model. J Clin Virol 52, 215–221 (2011).

2. Abed, Y. & Boivin, G. Treatment of respiratory virus infections. Antiviral Res 70, 1–16 (2006).

3. Brandtzaeg, P. Function of Mucosa-Associated Lymphoid Tissue in Antibody Formation. Immunological Investigations 39, 303–355 (2010).

4. Holmgren, J. & Czerkinsky, C. Mucosal immunity and vaccines. Nature Medicine 11, S45–S53 (2005).

5. Yusuf, H. & Kett, V. Current prospects and future challenges for nasal vaccine delivery. Hum Vaccin Immunother 13, 34–45 (2017).

6. Lycke, N. Recent progress in mucosal vaccine development: potential and limitations. Nature Reviews Immunology 12, 592–605 (2012).

7. Zhou, S. et al. MyD88 is critical for the development of innate and adaptive immunity during acute lymphocytic choriomeningitis virus infection. Eur J Immunol 35, 822–830 (2005).

8. Dubey, V. & MacFadden, D. Disseminated varicella zoster virus infection after vaccination with a live attenuated vaccine. Canadian Medical Association Journal 191, E1025–E1027 (2019).

9. Danziger-Isakov, L., Kumar, D. & Practice, T.A.I.C.o. Vaccination of solid organ transplant candidates and recipients: Guidelines from the American society of transplantation infectious diseases community of practice. Clinical Transplantation 33, e13563 (2019).

10. Sanders, B., Koldijk, M. & Schuitemaker, H. Inactivated Viral Vaccines. Vaccine Analysis: Strategies, Principles, and Control, 45–80 (2014).

11. Sumner, R.P., Ren, H., Ferguson, B.J. & Smith, G.L. Increased attenuation but decreased immunogenicity by deletion of multiple vaccinia virus immunomodulators. Vaccine 34, 4827–4834 (2016).

12. Moyle, P.M. & Toth, I. Modern Subunit Vaccines: Development, Components, and Research Opportunities. ChemMedChem 8, 360–376 (2013).

13. Pulendran, B. & Ahmed, R. Immunological mechanisms of vaccination. Nat Immunol 12, 509–517 (2011).

14. Pulendran, B., S. Arunachalam, P. & O’Hagan, D.T. Emerging concepts in the science of vaccine adjuvants. Nature Reviews Drug Discovery (2021).

15. Wang, Z.-B. & Xu, J. Better Adjuvants for Better Vaccines: Progress in Adjuvant Delivery Systems, Modifications, and Adjuvant-Antigen Codelivery. Vaccines (Basel) 8, 128 (2020).

16. McGhee, J.R. & Fujihashi, K. Inside the Mucosal Immune System. PLOS Biology 10, e1001397 (2012).

17. Freytag, L.C. & Clements, J.D. Chapter 61 - Mucosal Adjuvants: New Developments and Challenges. In: Mestecky, J., et al. (eds). Mucosal Immunology (Fourth Edition). Academic Press: Boston, 2015, pp 1183-1199.

18. Li, S. et al. Type I Interferons: Distinct Biological Activities and Current Applications for Viral Infection. Cellular Physiology and Biochemistry 51, 2377–2396 (2018).

19. Su, F., Patel, G.B., Hu, S. & Chen, W. Induction of mucosal immunity through systemic immunization: Phantom or reality? Hum Vaccin Immunother 12, 1070–1079 (2016).

20. Vassilieva, E.V., Taylor, D.W. & Compans, R.W. Combination of STING Pathway Agonist With Saponin Is an Effective Adjuvant in Immunosenescent Mice. Frontiers in Immunology 10 (2019).

21. Van Dis, E. et al. STING-Activating Adjuvants Elicit a Th17 Immune Response and Protect against Mycobacterium tuberculosis Infection. Cell Rep 23, 1435–1447 (2018).

22. Wang, H. et al. cGAS is essential for the antitumor effect of immune checkpoint blockade. Proceedings of the National Academy of Sciences of the United States of America 114, 1637–1642 (2017).

23. Osorio, F. & e Sousa, C.R. Myeloid C-type lectin receptors in pathogen recognition and host defense. Immunity 34, 651–664 (2011).

24. Zaharoff, D.A., Rogers, C.J., Hance, K.W., Schlom, J. & Greiner, J.W. Chitosan solution enhances both humoral and cell-mediated immune responses to subcutaneous vaccination. Vaccine 25, 2085–2094 (2007).

25. Carroll, E.C. et al. The vaccine adjuvant chitosan promotes cellular immunity via DNA sensor cGAS-STING-dependent induction of type I interferons. Immunity 44, 597–608 (2016).

26. Hoover, A.R., et al. A novel biopolymer synergizes type I IFN and IL-1β production through STING. bioRxiv, 2022.2007.2022.501157 (2022).

27. Li, X., et al. Chitin, Chitosan, and Glycated Chitosan Regulate Immune Responses: The Novel Adjuvants for Cancer Vaccine. Clinical and Developmental Immunology 2013, 387023 (2013).

28. Zhao, D. et al. Biomedical Applications of Chitosan and Its Derivative Nanoparticles. Polymers (Basel) 10, 462 (2018).

29. Bellich, B., D’Agostino, I., Semeraro, S., Gamini, A. & Cesàro, A. “The Good, the Bad and the Ugly” of Chitosans. Marine Drugs 14, 99 (2016).

30. Li, X. et al. Preliminary safety and efficacy results of laser immunotherapy for the treatment of metastatic breast cancer patients. Photochem Photobiol Sci 10, 817–821 (2011).

31. Liu, K. et al. Antigen presentation and interferon signatures in B cells driven by localized ablative cancer immunotherapy correlate with extended survival. Theranostics 12, 639–656 (2022).

32. Korbelik, M., Hode, T., Lam, S.S. & Chen, W.R. Novel Immune Stimulant Amplifies Direct Tumoricidal Effect of Cancer Ablation Therapies and Their Systemic Antitumor Immune Efficacy. Cells 10, 492 (2021).

33. Barchfeld, G.L. et al. The adjuvants MF59 and LT-K63 enhance the mucosal and systemic immunogenicity of subunit influenza vaccine administered intranasally in mice. Vaccine 17, 695–704 (1999).

34. Cubuk, J. et al. The SARS-CoV-2 nucleocapsid protein is dynamic, disordered, and phase separates with RNA. Nature Communications 12, 1936 (2021).

35. Dai, Y. et al. Immunodominant regions prediction of nucleocapsid protein for SARS-CoV-2 early diagnosis: a bioinformatics and immunoinformatics study. Pathog Glob Health 114, 463–470 (2020).

36. Lavelle, E.C. & Ward, R.W. Mucosal vaccines — fortifying the frontiers. Nature Reviews Immunology 22, 236–250 (2022).

37. Reed, S.G., Orr, M.T. & Fox, C.B. Key roles of adjuvants in modern vaccines. Nature Medicine 19, 1597–1608 (2013).

38. Stetson, D.B. & Medzhitov, R. Type I interferons in host defense. Immunity 25, 373–381 (2006).

39. Fink, K. et al. Early type[]I interferon-mediated signals on B cells specifically enhance antiviral humoral responses. Eur J Immunol 36, 2094–2105 (2006).

40. Kolumam, G.A., Thomas, S., Thompson, L.J., Sprent, J. & Murali-Krishna, K. Type I interferons act directly on CD8 T cells to allow clonal expansion and memory formation in response to viral infection. The Journal of experimental medicine 202, 637–650 (2005).

41. Kuo, T.-Y. et al. Development of CpG-adjuvanted stable prefusion SARS-CoV-2 spike antigen as a subunit vaccine against COVID-19. Scientific Reports 10, 20085 (2020).

42. Campbell, J.D. Development of the CpG Adjuvant 1018: A Case Study. Methods in molecular biology (Clifton, N.J.) 1494, 15–27 (2017).

43. Hoover, A.R., et al. ScRNA-seq reveals tumor microenvironment remodeling induced by local intervention-based immunotherapy. bioRxiv, 2020.2010.2002.323006 (2020).

44. Sallusto, F. & Lanzavecchia, A. The instructive role of dendritic cells on T-cell responses. Arthritis Res 4 Suppl 3, S127–S132 (2002).

45. Hoover, A.R. et al. Single-cell RNA sequencing reveals localized tumour ablation and intratumoural immunostimulant delivery potentiate T cell mediated tumour killing. Clinical and Translational Medicine 12, e937 (2022).

46. Zhou, F. et al. Local phototherapy synergizes with immunoadjuvant for treatment of pancreatic cancer through induced immunogenic tumor vaccine. Clin. Cancer Res. 24, 5335–5346 (2018).

47. El-Hussein, A., Lam, S.S., Raker, J., Chen, W.R. & Hamblin, M.R. N-Dihydrogalactochitosan as a potent immune activator for dendritic cells. Journal of Biomedical Materials Research Part A (2016).

48. Curtsinger, J.M., Valenzuela, J.O., Agarwal, P., Lins, D. & Mescher, M.F. Cutting Edge: Type I IFNs Provide a Third Signal to CD8 T Cells to Stimulate Clonal Expansion and Differentiation. The Journal of Immunology 174, 4465–4469 (2005).

49. Ben-Sasson, S.Z., Wang, K., Cohen, J. & Paul, W.E. IL-1β strikingly enhances antigen-driven CD4 and CD8 T-cell responses. Cold Spring Harbor symposia on quantitative biology 78, 117–124 (2013).

50. Oliveira, S.C., de Magalhães, M.T.Q. & Homan, E.J. Immunoinformatic Analysis of SARS-CoV-2 Nucleocapsid Protein and Identification of COVID-19 Vaccine Targets. Frontiers in Immunology 11 (2020).

51. Smits, V.A.J. et al. The Nucleocapsid protein triggers the main humoral immune response in COVID-19 patients. Biochemical and Biophysical Research Communications 543, 45–49 (2021).

52. Lee, E. et al. Identification of SARS-CoV-2 Nucleocapsid and Spike T-Cell Epitopes for Assessing T-Cell Immunity. J Virol 95 (2021).

53. Lineburg, K.E. et al. CD8(+) T cells specific for an immunodominant SARS-CoV-2 nucleocapsid epitope cross-react with selective seasonal coronaviruses. Immunity 54, 1055–1065.e1055 (2021).

54. Carotti, M. et al. Chest CT features of coronavirus disease 2019 (COVID-19) pneumonia: key points for radiologists. La radiologia medica 125, 636–646 (2020).

55. Tsang, K.W. & Bilton, D. Clinical challenges in managing bronchiectasis. Respirology 14, 637–650 (2009).

56. Ambrosetti, M.C. et al. Rapid onset of bronchiectasis in COVID-19 Pneumonia: two cases studied with CT. Radiol Case Rep 15, 2098–2103 (2020).

57. Wilkins, A.L. et al. AS03- and MF59-Adjuvanted Influenza Vaccines in Children. Frontiers in Immunology 8 (2017).

58. McCray, P.B., Jr. et al. Lethal infection of K18-hACE2 mice infected with severe acute respiratory syndrome coronavirus. J Virol 81, 813–821 (2007).

