## Supplementary material for "A novel biopolymer for mucosal adjuvant against respiratory pathogens": suppl figure legends

**Supplemental Figure 1. GC retains antigen at the site of injection and stimulates a localized immune response.**

A. Schematic of the injection and imaging protocol to visualize antigen retention.

B. Retention of antigen by GC. GC (0.9, 0.75, or 0.5%) was mixed with ovalbumin labeled by Texas-Red (OVA-TR) and injected intradermally into the flank of C57BL/6 mice. Whole-body fluorescent imaging was performed over a period of several days. The percentage rate of change was calculated to quantify the amount of ovalbumin localized at the injection site. Two-way ANOVA was used for statistical analysis.

C. Activation of leukocytes by GC. Mice received an intranasal application of 1x PBS or 1% GC (20μl per nostril) and 72 hours later cervical lymph nodes were isolated and flow cytometry was used to analyze the leukocytes. B cells were gated using Live, CD45^+^CD19^+^B220^+^, migrating DCs were gated using CD45^+^CD11^High^MHC-II^High^, T cells were gated using CD45^+^CD3^+^ and either CD4^+^ or CD8^+^, and resident DCs were gated using CD45^+^CD11c^int^MHC-II^int^. This gating strategy was used to calculate the cell populations. Unpaired t-test was used for statistical analysis.

D. Stimulation of antiviral response pathways by GC. Wild-type BMDCs were stimulated +/- GC for 24 hours prior to harvesting for bulk RNA-sequencing analysis. Gene Ontology Pathways Analysis revealed that GC stimulates a variety of antiviral response pathways.

**Supplemental Figure 2.** **One-time (1X) intranasal vaccination (Vax) using GC with recombinant viral proteins (RVPs) produces minimal changes two weeks post-vaccination.**

A. Schematic of the one vaccination (1X) strategy and analysis.

B. Total serum IgG and IgA levels two weeks after one vaccination. One-way ANOVA was used for statistical analysis.

C-E. Activation of leukocytes after one vaccination. Cervical lymph nodes were isolated two weeks after the intranasal vaccination and analyzed via flow cytometry.

C. Activation of CD8^+^ T cells after one vaccination. CD8^+^ T cells were gated on Live, CD45^+^CD3^+^CD8^+^ cells prior to analysis with CD44 and CD62L. Total cellularity of CD8^+^ T cells, CD8^+^CD44^+^, CD8^+^CD44^+^CD62L^+^, and CD8^+^CD62L^+^ was calculated. One-way ANOVA was used for statistical analysis.

D. Activation of CD4^+^ T cells after one vaccination. CD4^+^ T cells were gated on Live, CD45^+^CD3^+^CD4^+^ cells prior to analysis with CD44 and CD62L. Total cellularity of CD4^+^ T cells, CD4^+^CD44^+^, CD4^+^CD44^+^CD62L^+^, and CD4^+^CD62L^+^ was calculated. One-way ANOVA was used for statistical analysis.

E. Activation of B cells after one vaccination. B cells were gated on Live, CD45^+^CD19^+^B220^+^ cells and then calculated for total cellularity. One-way ANOVA was used for statistical analysis.

**Supplemental Figure 3. Intranasal vaccination of GC combined with recombinant SARS-CoV-2 spike protein results in minimal leukocyte alterations in the lungs.**

A-C. Effects of GC on leukocytes in the lungs. The animals were vaccinated according to the schematic in Figure 2A. Lungs were harvested two weeks post-second vaccination. Lungs were isolated, single-cell suspensions were generated, and leukocytes were analyzed by flow cytometry.

A. Activation of CD8^+^ T cells two weeks after the second vaccination. CD8^+^ T cells were gated on Live, CD45^+^CD3^+^CD8^+^ cells prior to analysis with CD44 and CD62L. Total cellularity of CD8 T cells, CD8^+^CD44^+^, CD8^+^CD44^+^CD62L^+^, and CD8^+^CD62L^+^ was calculated. One-way ANOVA was used for statistical analysis.

B. Activation of CD4^+^ T cells two weeks after the second vaccination. CD4^+^ T cells were gated on Live, CD45^+^CD3^+^CD4^+^ cells prior to analysis with CD44 and CD62L. Total cellularity of CD4 T cells, CD4^+^CD44^+^, CD4^+^CD44^+^CD62L^+^, and CD4^+^CD62L^+^ was calculated. One-way ANOVA was used for statistical analysis.

C. Activation of B cells two weeks after the second vaccination. B cells were gated on Live, CD45^+^CD19^+^B220^+^ cells and then calculated for total cellularity. One-way ANOVA was used for statistical analysis.

**Supplemental Figure 4. Intranasal vaccination of GC combined with recombinant SARS-CoV-2 spike and nucleocapsid proteins results in elevated leukocyte numbers in the lungs.**

A-C. Activation of leukocytes in the lungs by GC with recombinant proteins. The animals were vaccinated according to the schematic in Figure 2A. Lungs were harvested two weeks after the second vaccination. Lungs were isolated, single-cell suspensions were generated, and leukocytes were analyzed by flow cytometry.

A. CD4^+^ T cells were gated on Live, CD45^+^CD3^+^CD4^+^ cells prior to analysis with CD44 and CD62L. Total cellularity of CD4 T cells, CD4^+^CD44^+^, CD4^+^CD44^+^CD62L^+^, and CD4^+^CD62L^+^ was calculated. One-way ANOVA was used for statistical analysis.

B. CD8^+^ T cells were gated on Live, CD45^+^CD3^+^CD8^+^ cells prior to analysis with CD44 and CD62L. Total cellularity of CD8 T cells, CD8^+^CD44^+^, CD8^+^CD44^+^CD62L^+^, and CD8^+^CD62L^+^ was calculated. One-way ANOVA was used for statistical analysis.

C. B cells were gated on Live, CD45^+^CD19^+^B220^+^ cells and then calculated for total cellularity. One-way ANOVA was used for statistical analysis.

**Supplemental Figure 5. Subcutaneous vaccination with GC or AV and recombinant SARS-CoV-2 spike and nucleocapsid proteins protects mice against lethal viral challenge.**

A. Schematic of the sub-Q vaccination and viral challenge strategy for K18-hACE-2 animals.

B. Survival rates of animals challenged by SARS-CoV-2 virus over a course of 21 days (n=6).

C. Percent body weight change over a period of 21 days in virally infected animals.
