## Supplementary material for "A novel biopolymer for mucosal adjuvant against respiratory pathogens": suppl figures

### Slide 1
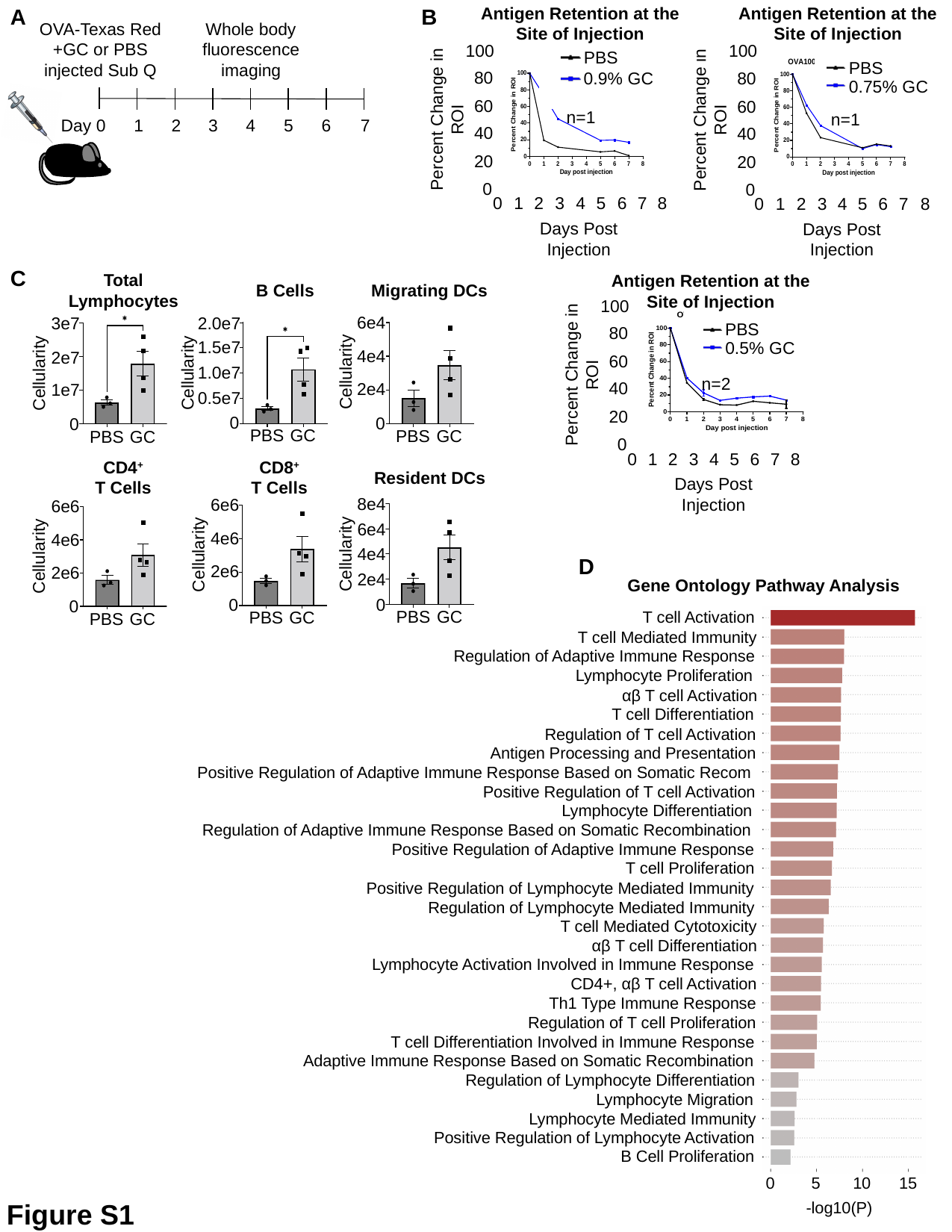

Antigen Retention at the Site of Injection
Antigen Retention at the Site of Injection
A
B
OVA-Texas Red +GC or PBS injected Sub Q
Whole body fluorescence imaging
Day 0
1
2
3
4
5
6
7
100
80
60
Percent Change in ROI
40
20
0
0
1
2
3
4
5
6
7
Days Post Injection
8
PBS
0.75% GC
n=1
100
80
60
Percent Change in ROI
40
20
0
0
1
2
3
4
5
6
7
Days Post Injection
8
PBS
0.9% GC
n=1
C
Total
Lymphocytes
3e7
2e7
Cellularity
1e7
0
PBS
GC
B Cells
2.0e7
1.5e7
1.0e7
Cellularity
0.5e7
0
PBS
GC
Migrating DCs
6e4
4e4
Cellularity
2e4
0
PBS
GC
CD4^+^
T Cells
CD8^+^
T Cells
Resident DCs
8e4
6e6
4e6
Cellularity
2e6
0
PBS
GC
6e6
4e6
Cellularity
2e6
0
PBS
GC
6e4
4e4
Cellularity
2e4
0
PBS
GC
Antigen Retention at the Site of Injection
100
80
60
Percent Change in ROI
40
20
0
0
1
2
3
4
5
6
7
Days Post Injection
PBS
0.5% GC
n=2
8
D
Gene Ontology Pathway Analysis
T cell Activation
D
T cell Mediated Immunity
Regulation of Adaptive Immune Response
Lymphocyte Proliferation
αβ T cell Activation
T cell Differentiation
Regulation of T cell Activation
Antigen Processing and Presentation
Positive Regulation of Adaptive Immune Response Based on Somatic Recom
Positive Regulation of T cell Activation
Lymphocyte Differentiation
Regulation of Adaptive Immune Response Based on Somatic Recombination
Positive Regulation of Adaptive Immune Response
T cell Proliferation
Positive Regulation of Lymphocyte Mediated Immunity
Regulation of Lymphocyte Mediated Immunity
T cell Mediated Cytotoxicity
αβ T cell Differentiation
Lymphocyte Activation Involved in Immune Response
CD4+, αβ T cell Activation
Th1 Type Immune Response
Regulation of T cell Proliferation
T cell Differentiation Involved in Immune Response
Adaptive Immune Response Based on Somatic Recombination
Regulation of Lymphocyte Differentiation
Lymphocyte Migration
Lymphocyte Mediated Immunity
Positive Regulation of Lymphocyte Activation
B Cell Proliferation
0
5
10
15
-log10(P)
Figure S1

### Slide 2
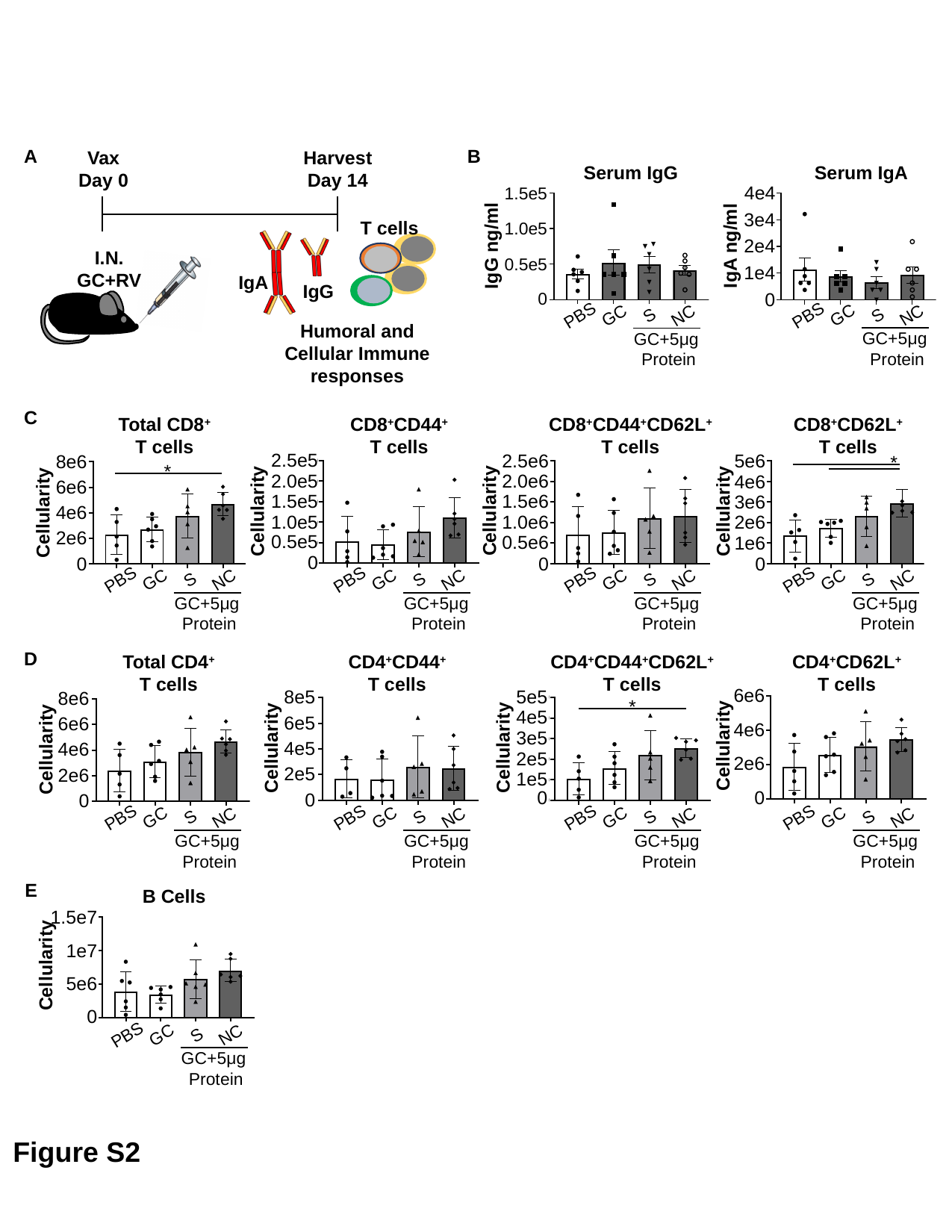

A
B
Vax
Day 0
Harvest
Day 14
Serum IgG
Serum IgA
4e4
1.5e5
3e4
2e4
IgA ng/ml
1e4
0
T cells
1.0e5
IgG ng/ml
I.N. GC+RVPs
0.5e5
IgA
IgG
0
PBS
GC
S
NC
GC+5μg Protein
PBS
GC
S
NC
GC+5μg Protein
Humoral and Cellular Immune responses
C
Total CD8^+^
T cells
CD8^+^CD44^+^
T cells
CD8^+^CD44^+^CD62L^+^
T cells
CD8^+^CD62L^+^
T cells
2.5e5
5e6
2.5e6
8e6
*
*
2.0e5
2.0e6
4e6
6e6
1.5e5
1.5e6
3e6
Cellularity
Cellularity
Cellularity
4e6
Cellularity
1.0e5
2e6
1.0e6
2e6
0.5e5
0.5e6
1e6
0
0
0
0
PBS
GC
S
NC
GC+5μg Protein
PBS
GC
S
NC
GC+5μg Protein
PBS
GC
S
NC
GC+5μg Protein
PBS
GC
S
NC
GC+5μg Protein
D
Total CD4^+^
T cells
CD4^+^CD44^+^
T cells
CD4^+^CD44^+^CD62L^+^
T cells
CD4^+^CD62L^+^
T cells
6e6
5e5
8e5
8e6
*
4e5
6e5
6e6
4e6
3e5
Cellularity
4e5
Cellularity
Cellularity
4e6
Cellularity
2e5
2e6
2e5
2e6
1e5
0
0
0
0
PBS
GC
S
NC
GC+5μg Protein
PBS
GC
S
NC
GC+5μg Protein
PBS
GC
S
NC
GC+5μg Protein
PBS
GC
S
NC
GC+5μg Protein
E
B Cells
1.5e7
1e7
Cellularity
5e6
0
PBS
GC
S
NC
GC+5μg Protein
Figure S2

### Slide 3
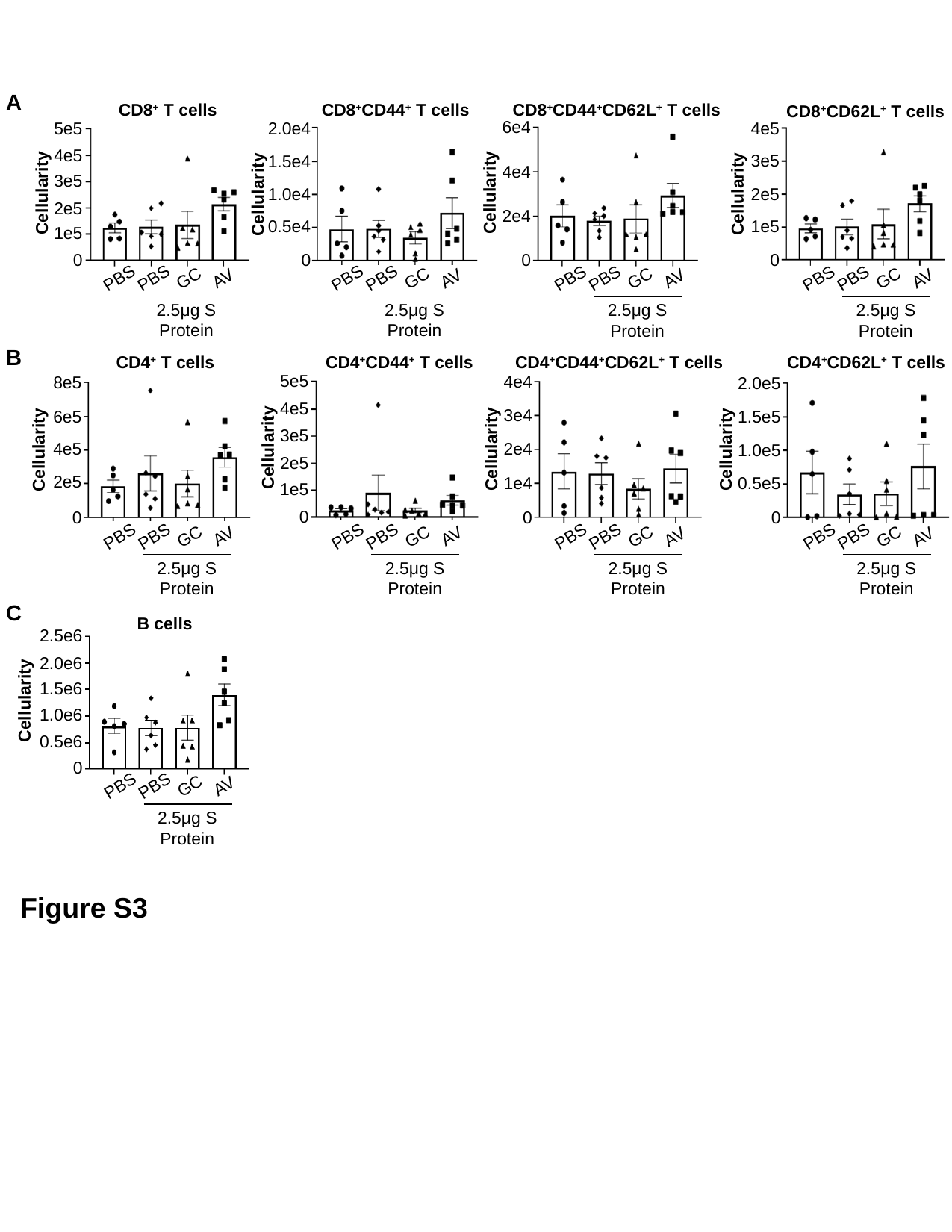

A
CD8^+^ T cells
CD8^+^CD44^+^ T cells
CD8^+^CD44^+^CD62L^+^ T cells
CD8^+^CD62L^+^ T cells
6e4
4e5
5e5
2.0e4
4e5
3e5
1.5e4
4e4
3e5
Cellularity
Cellularity
1.0e4
Cellularity
2e5
Cellularity
2e5
2e4
1e5
0.5e4
1e5
0
0
0
0
PBS
PBS
GC
AV
2.5μg S Protein
PBS
PBS
GC
AV
2.5μg S Protein
PBS
PBS
GC
AV
2.5μg S Protein
PBS
PBS
GC
AV
2.5μg S Protein
B
CD4^+^ T cells
CD4^+^CD44^+^ T cells
CD4^+^CD44^+^CD62L^+^ T cells
CD4^+^CD62L^+^ T cells
5e5
4e4
8e5
2.0e5
4e5
3e4
6e5
1.5e5
3e5
Cellularity
4e5
2e4
Cellularity
Cellularity
Cellularity
1.0e5
2e5
2e5
0.5e5
1e4
1e5
0
0
0
0
PBS
PBS
GC
AV
2.5μg S Protein
PBS
PBS
GC
AV
2.5μg S Protein
PBS
PBS
GC
AV
2.5μg S Protein
PBS
PBS
GC
AV
2.5μg S Protein
C
B cells
2.5e6
2.0e6
1.5e6
Cellularity
1.0e6
0.5e6
0
PBS
PBS
GC
AV
2.5μg S Protein
Figure S3

### Slide 4
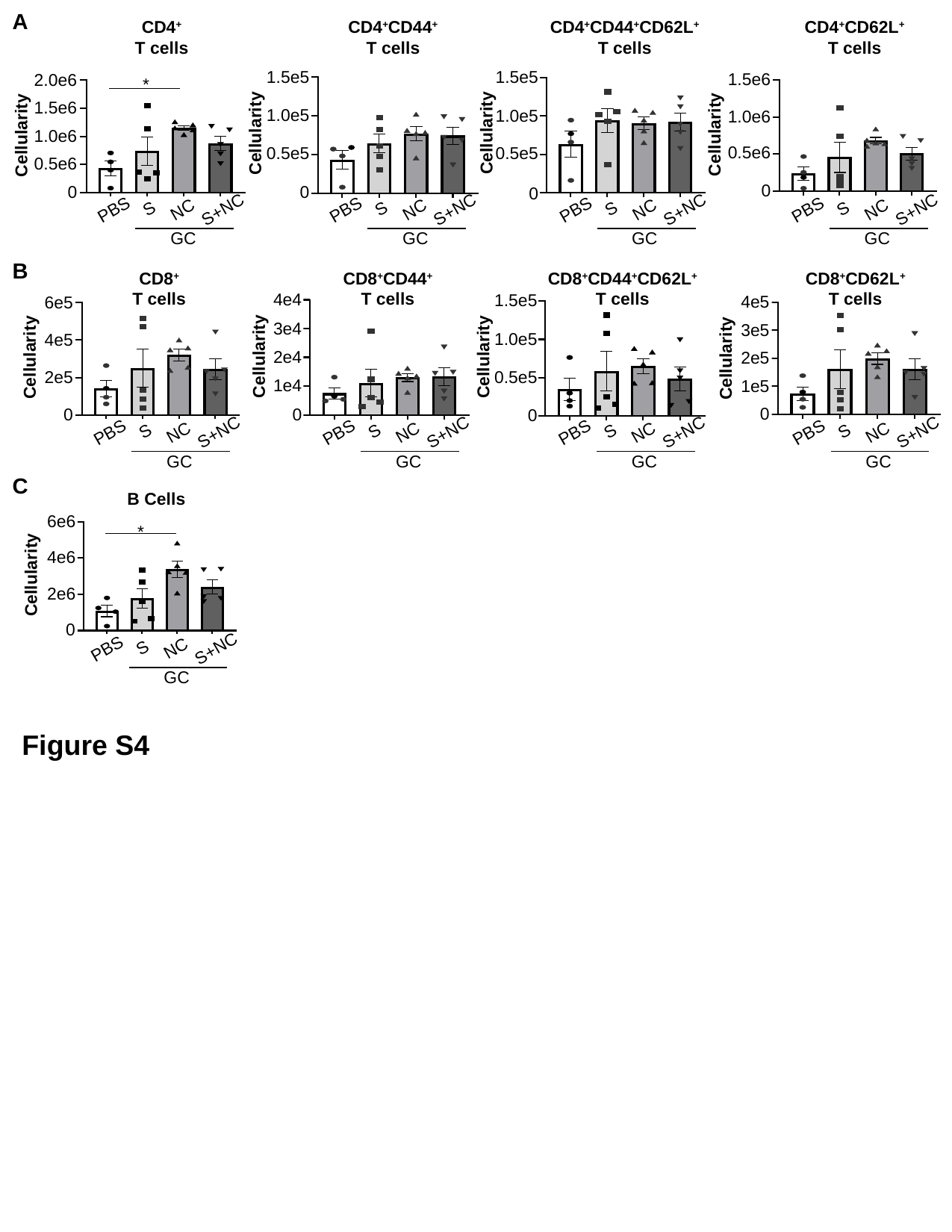

A
CD4^+^
T cells
CD4^+^CD44^+^
T cells
CD4^+^CD44^+^CD62L^+^
T cells
CD4^+^CD62L^+^
T cells
1.5e5
1.5e5
1.5e6
2.0e6
*
1.5e6
1.0e5
1.0e5
1.0e6
Cellularity
Cellularity
Cellularity
Cellularity
1.0e6
0.5e6
0.5e5
0.5e5
0.5e6
0
0
0
0
PBS
S
NC
S+NC
GC
PBS
S
NC
S+NC
GC
PBS
S
NC
S+NC
GC
PBS
S
NC
S+NC
GC
B
CD8^+^
T cells
CD8^+^CD44^+^
T cells
CD8^+^CD44^+^CD62L^+^
T cells
CD8^+^CD62L^+^
T cells
4e4
1.5e5
4e5
6e5
3e4
3e5
1.0e5
4e5
2e4
Cellularity
Cellularity
2e5
Cellularity
Cellularity
2e5
0.5e5
1e4
1e5
0
0
0
0
PBS
S
NC
S+NC
GC
PBS
S
NC
S+NC
GC
PBS
S
NC
S+NC
GC
PBS
S
NC
S+NC
GC
C
B Cells
6e6
*
4e6
Cellularity
2e6
0
PBS
S
NC
S+NC
GC
Figure S4

### Slide 5
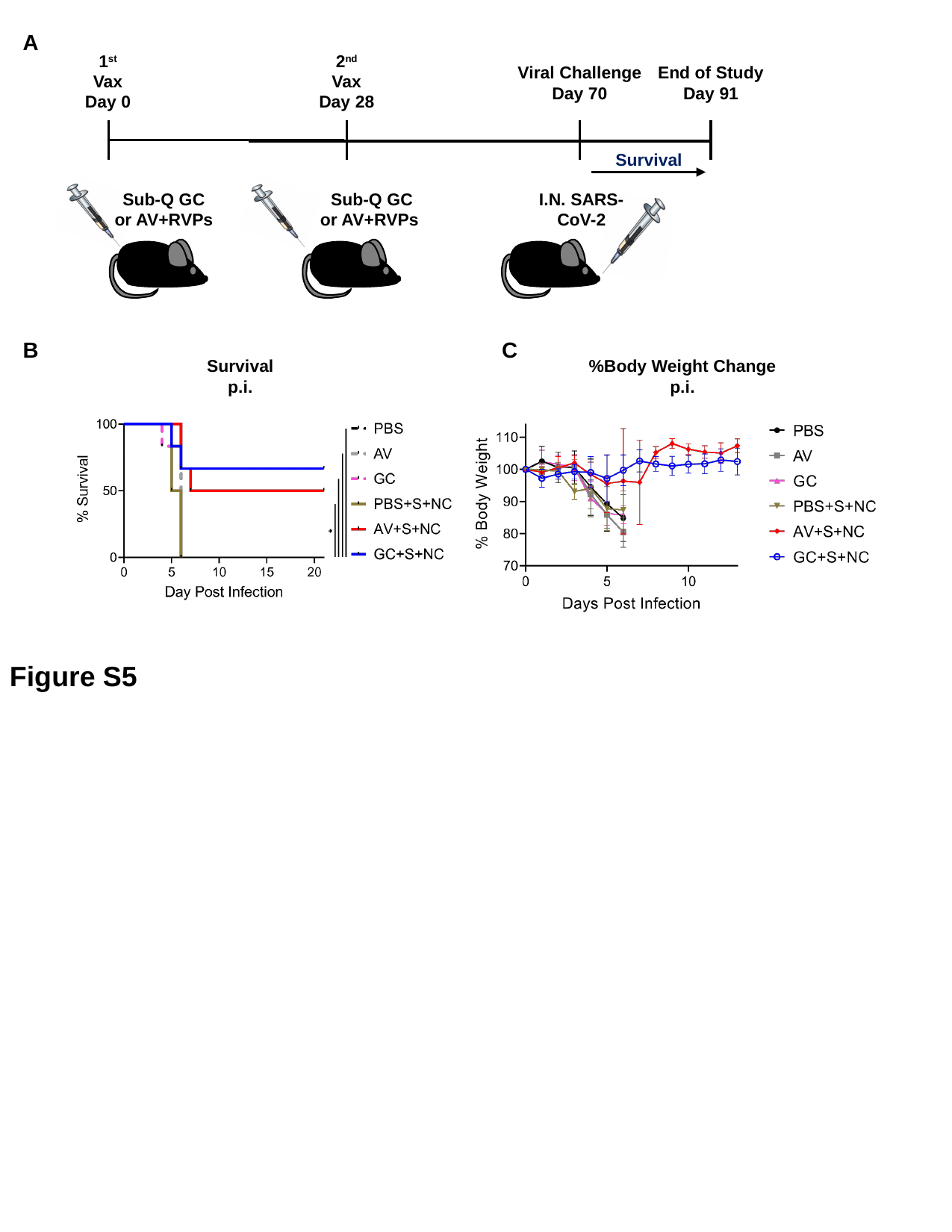

A
1^st^
Vax
Day 0
2^nd^
Vax
Day 28
Viral Challenge
Day 70
End of Study
Day 91
Survival
Sub-Q GC or AV+RVPs
Sub-Q GC or AV+RVPs
I.N. SARS-CoV-2
B
C
Survival
p.i.
%Body Weight Change
p.i.
Figure S5
